# Temozolomide-induced guanine mutations create exploitable vulnerabilities of guanine-rich DNA and RNA regions in drug resistant gliomas

**DOI:** 10.1101/661660

**Authors:** Deanna M Tiek, Beril Erdogdu, Roham Razaghi, Lu Jin, Norah Sadowski, Carla Alamillo-Ferrer, J Robert Hogg, Bassem R Haddad, David H Drewry, Carrow I Wells, Julie E. Pickett, Xiao Song, Anshika Goenka, Bo Hu, Samuel L Goldlust, William J Zuercher, Mihaela Pertea, Winston Timp, Shi-Yuan Cheng, Rebecca B Riggins

## Abstract

Temozolomide (TMZ) is a chemotherapeutic agent that has been the first-line standard of care for the aggressive brain cancer glioblastoma (GBM) since 2005. Though initially beneficial, TMZ-resistance is universal and second-line interventions are an unmet clinical need. Here we took advantage the mechanism of action of TMZ to target guanines (G) and investigated G-rich g-quadruplex (G4) and splice site changes that occur upon TMZ-resistance. We report TMZ-resistant GBM has guanine mutations that disrupt the G-rich DNA G4s and splice sites that lead to deregulated alternative splicing. These alterations create vulnerabilities, which are selectively targeted by either the G4 stabilizing drug TMPyP4 or a novel splicing kinase inhibitor of cdc2- like kinase. Finally, we show that the G4 and RNA-binding protein EWSR1 aggregates in the cytoplasm in TMZ-resistant GBM cells and patient samples. Together, our findings provide insight into targetable vulnerabilities of TMZ-resistant GBM and present cytoplasmic EWSR1 as a putative biomarker.

**Teaser:** Targeting temozolomide mutations in drug resistant glioma via g-quadruplex and splicing modulators with a putative biomarker.

## Introduction

GBM is the most diagnosed glioma and has an abysmal overall survival rate of ∼5% at 5 years (1). With a median survival time of 14∼16 months, it is uniformly fatal (2). Temozolomide (TMZ; Temodar) is the FDA-approved standard of care first line therapy for GBM in combination with surgery and radiation (3). TMZ causes DNA damage by adding mutagenic adducts to DNA, predominantly O^6^-methyl guanine, a lesion that can be repaired by the suicide DNA repair protein methyl guanine methyl transferase (MGMT) (4). The overall survival benefit provided by TMZ is ∼ 4 months, and rapid development of TMZ resistance occurs at least in part through demethylation of the MGMT promoter, allowing for the expression of MGMT (5).

TMZ preferentially targets guanines (6), critical nucleotides in many DNA and RNA secondary structures (7). Two structurally and functionally important and distinct G-rich regions are g-quadruplexes (G4s) and RNA splice sites. Previous studies have shown G4s to be important regulatory elements for oncogenes like *c-MYC*, *KIT*, or *KRAS* (7). These G4s have diverse structural and physicochemical properties that suggest a high degree of selectivity, and thus G4s may represent a selective druggable target in multiple cancers (8). In contrast, studies of *C9orf72* in amyotrophic lateral sclerosis (ALS), a neurodegenerative disease show that alterations of G4 tracts can cause nucleolar stress, deregulate RNA alternative splicing (AS), and alter RNA binding protein localization (9). Nucleolar stress, manifested as changes in nucleolar size and circularity, is a poor prognostic factor in many cancers, including GBM (10).

Changes in RNA alternative splicing (AS) patterns are widespread in GBM, with prior work suggesting that GBM is “addicted to splicing” (11). Previous work has shown that targeting splicing signatures may be effective in certain types of gliomas. Braun *et al* showed that gliomas have a high propensity to accumulate detained introns, which can be targeted through PRMT5 inhibition (11). Other splicing families, like the serine/arginine rich (SR) family and cdc2-like kinases (CLKs) have also been shown to play modulatory roles in specific oncogenic processes(12). One family with essential functions in RNA processing in cancer and neurodegenerative diseases (NDs) is the FET family of proteins, consisting of FUS, EWSR1, and TAF15 (13). The aggregation-prone properties of the FET proteins have established FUS as a prototype to study protein aggregation, and a biomarker for ALS and frontotemporal dementia (FTLD) where EWSR1 has similar aggregation-prone properties (14). FET proteins also are involved in DNA repair (15), can bind the G4 secondary structure of DNA and RNA, and play roles in AS isoform fate (16).

In this study we sought to use the well-known mechanism of action of TMZ to define targetable dependencies of TMZ-resistant GBM. We went on to exploit the TMZ-induced DNA and RNA changes in TMZ-resistant GBM with both known and novel small molecules targeting G4s and the activity of splicing regulatory kinases. Finally, we propose EWSR1 cytoplasmic aggregation as a putative biomarker for TMZ-resistant GBM where AS modulators may be successful second-line interventions.

## Results

### TMZ-induced guanine mutations

Temozolomide (TMZ; Temodar), the FDA-approved standard of care first-line therapy for glioblastoma (GBM), adds mutagenic adducts to guanine, the most prevalent being O^6^- methylguanine. To determine the role of guanine (G) mutations in TMZ-resistance we performed whole genome sequencing (WGS) on patient-derived GBM cell lines that are TMZ-sensitive (42WT), have acquired resistance to TMZ through an *in vitro* selection (42R) (17), or the established GBM intrinsically TMZ-resistant line T98. Acquired and intrinsic GBM TMZ-resistant cells had significantly increased single nucleotide variants relative to the TMZ-sensitive cells (Fig. 1A). Further analysis showed G>A, and C>T point mutations were the most enriched in TMZ-resistant cells (Fig. 1B). These results are consistent with the previously defined Signature 11 (increased C>T point mutations) that is enriched in GBM patient samples and has a probable association with TMZ treatment (18).

**Figure 1.**
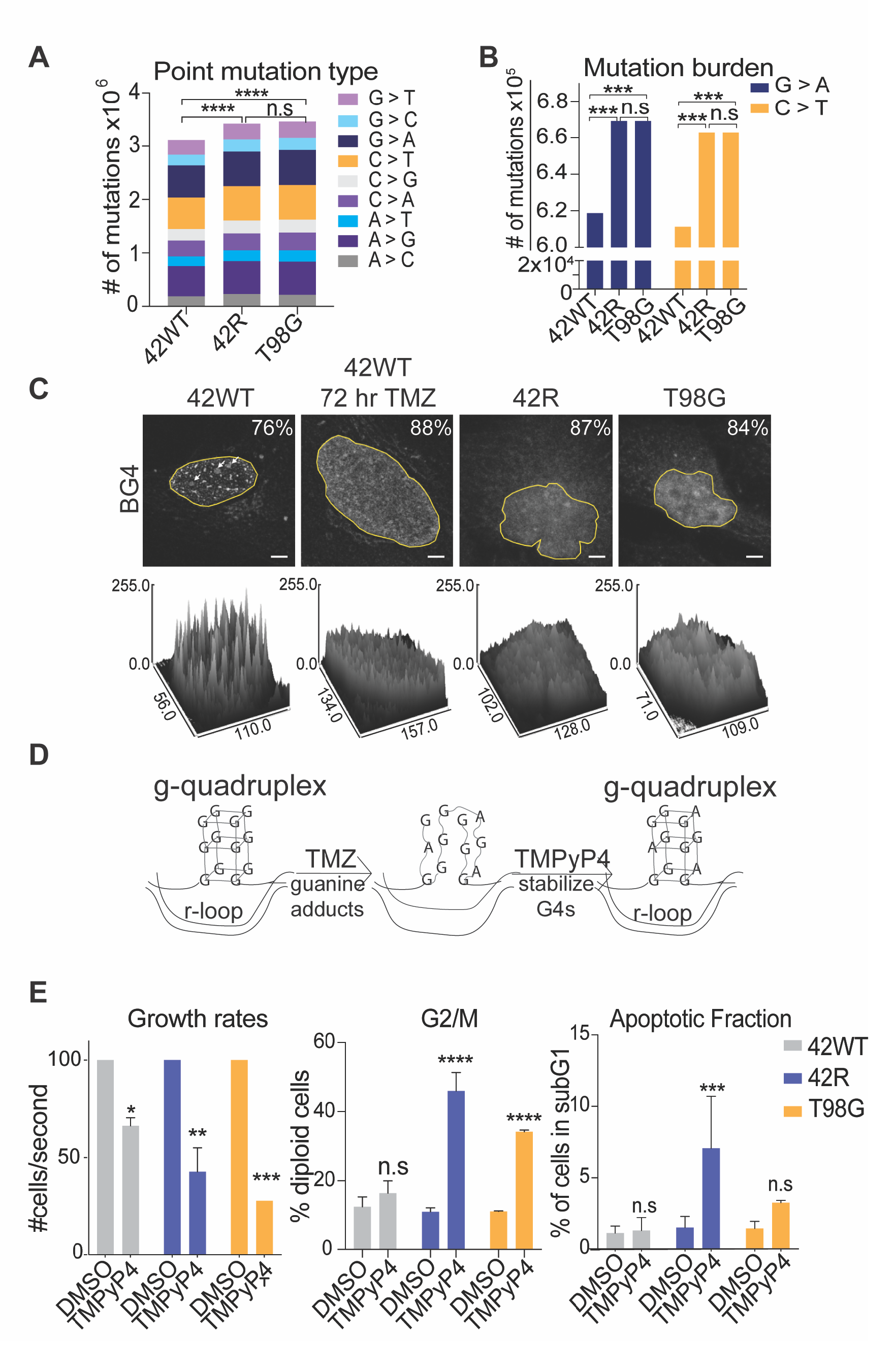
Guanine mutations in TMZ-resistant GBM cells disrupt G-quadruplex (G4) structure and associate with sensitivity to the G4 stabilizing drug TMPyP4. (A) Single nucleotide changes identified by whole genome sequencing data aligned to GRCh38 and analyzed by Genome Analysis Toolkit (GATK) mutation calling; duplicate samples with 2-way ANOVA for total mutation count. (B) Enrichment of G>A and C>T transition mutations in TMZ-resistant cells; duplicate samples with 2-way ANOVA as in (A). (C) Immunofluorescent staining (IF) with the G-quadruplex antibody BG4 in TMZ-sensitive 42WT, 72 hr-treated 42WT, TMZ-resistant 42R, and T98G cells with surface plots of signal intensity below generated by FIJI., Scale bar 5 µm. Yellow outline denotes the nucleus and the region analyzed in FIJI; representative of three biological replicates. (**D**) Graphical depiction of the hypothesis that TMZ-induced O6- methylguanine (O^6^meG) adducts and subsequent mutations disrupt G4s, and that the G4 stabilizing agent TMPyP4 may normalize G4 structure. (**E**) FACS analysis following 24 hours of treatment with 50 µM TMPyP4 or DMSO vehicle control in 42WT, 42R, and T98G cells; three biological replicates. Growth rates; t-test. Percent of cells in G2/M; t-test. Percent of cells with subG1 DNA content; t-test. *, p<0.05, **, p<0.001 *** p<0.0005, **** p<0.0001.

Two G-rich regions that are functionally and structurally important in cancer are G-quadruplexes (G4s) and RNA splice sites. G4s can be readily detected by bioinformatic analysis of WGS. To test the hypothesis that G4 structures would be perturbed in TMZ-resistant cells, we used Quadron, an algorithm that integrates sequence information to detect tracts of Gs and Cs with characteristic surrounding sequence features of G4s (19). After all G4s were detected, we identified G4 sequences unique to each cell line, and these unique G4s were then used to identify significantly enriched motifs using Multiple Em for Motif Elicitation (MEME) (20). In TMZ-sensitive 42WT cells the top three significantly enriched G-rich sequences contain solely G’s as hypothesized (Supplementary Fig. 1A). However, in the acquired resistant 42R cells we found enrichment of a sequence with higher representation of A and T’s (Supplementary Fig. 1A). The T98G line, which is intrinsically resistant and was not selected in culture with TMZ treatment, showed enrichment of C- and G/A-rich motifs (Supplementary Fig. 1A). These potential structural changes caused by mutations in G4s in the TMZ-resistant cells were then validated by immunofluorescence (IF) with the G4-detecting monoclonal antibody BG4. In the TMZ-sensitive 42WT cells, discrete nuclear puncta were detected by the BG4 antibody, in addition to more diffuse nucleolar staining (Fig. 1C). Confirming prior studies (21) showing BG4 specifically binds DNA G4s *in vivo*, treatment of 42WT cells with DNase I abrogated specific nuclear BG4 staining (Supplementary Fig. 1B). In the acute TMZ-treated 42WT cells and both TMZ-resistant cell lines, there were no longer discrete G4 nuclear puncta, which corroborated our WGS data (Fig. 1C). We then hypothesized that if disruption of G4 sequences played a functional role in TMZ-resistance, a G4-stabilizing drug may be more effective as a second line treatment in TMZ-resistant cells (Fig. 1D). Treatment with TMPyP4, a G4-stabilizing drug, caused a slight increase in the punctate BG4 staining pattern in 42R cells (Supplementary Fig. 1C). In both resistant cell lines, TMPyP4 treatment induced a significant G2/M arrest and decreased cell proliferation with increased sub G1 fraction only in the 42R resistant cell line, but had minimal effects on cell cycle profile and a slight decrease in growth in the TMZ sensitive cell line (Fig. 1E). These data support the hypothesis that G4-disrupting G mutations have a functional, and potentially targetable, role in TMZ-resistant GBM.

### Acquired nucleolar changes with TMZ-induced mutations in G-quadruplexes

G4s are abundant in nucleoli, major stress organelles in cells undergoing DNA damage, where nucleoli morphology is a poor prognostic marker in several cancers, including GBM (10). Nucleoli form around nuclear organizer regions (NOR) located on the p arm of the five acrocentric chromosomes: 13, 14, 15, 21, and 22. Analysis of metaphase spreads from 42WT vs. 42R cell lines showed an increase in total chromosome number in the 42R cell line (22). We reanalyzed these data to determine if the acrocentric, or nucleolar-forming, chromosomes were specifically hyper-enriched in the acquired resistant cell line and showed the 42R cells had a significantly higher fraction of acrocentric chromosomes compared to their TMZ-sensitive parental line (Fig. 2A). Next, we adapted the index of nucleolar disruption (iNo) score as a method of quantifying nucleolar stress that accounts for both size and roundness of nucleoli (10). Using immunofluorescent staining (IF) and automated analysis of the nucleolar protein nucleolin (NCL) to determine nucleolar size and circularity (Fig. 2B), we found significant increases in nucleolar size associated with both acute TMZ treatment (42WT+TMZ) and intrinsic resistance (T98G, Fig. 2C), and significant decreases in circularity, or nucleolar roundness, in acute TMZ-treated and both acquired and intrinsic TMZ-resistant nucleoli (Fig. 2D).

**Figure 2.**
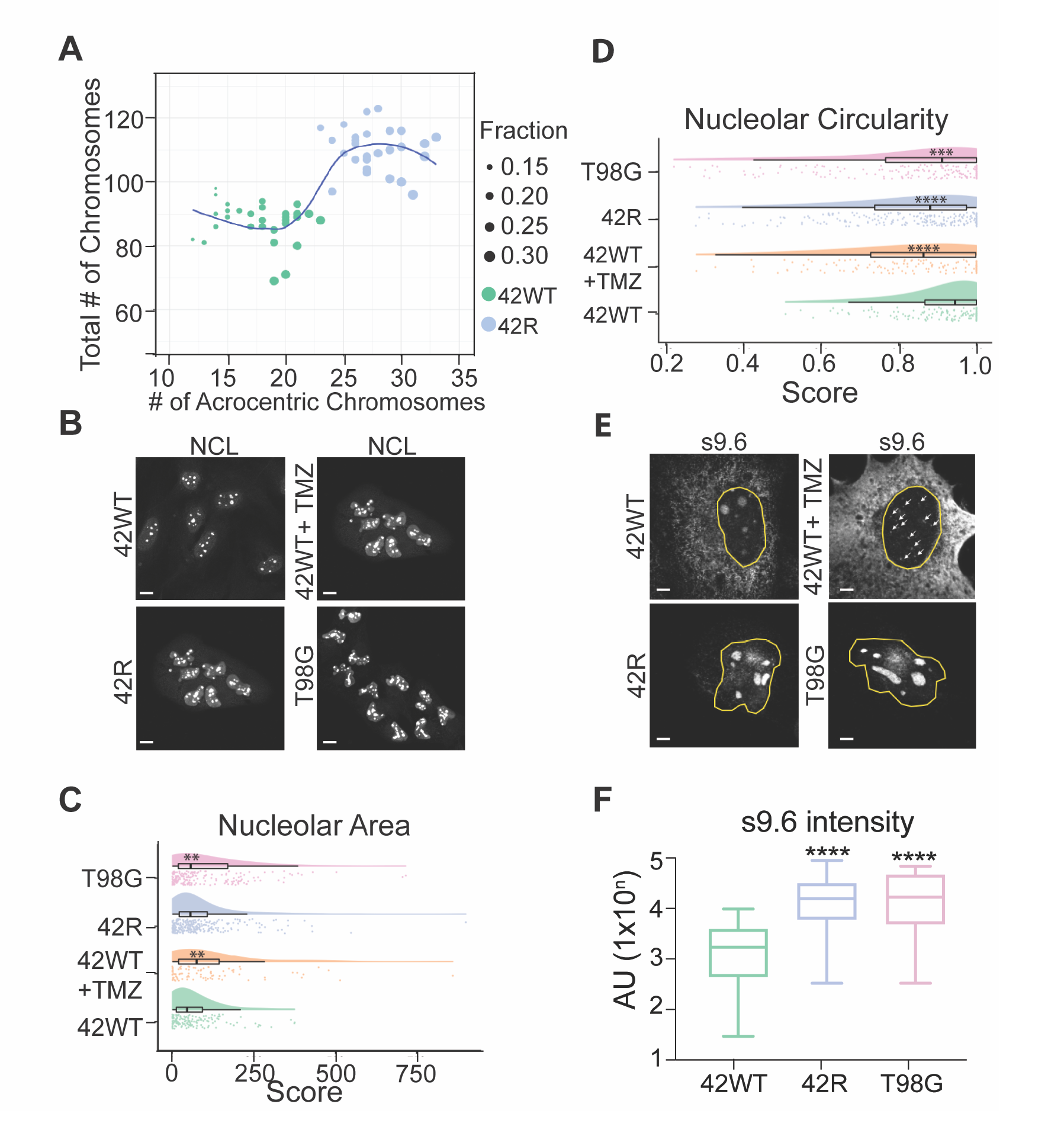
TMZ treatment and resistance associate with increased nucleolar stress and the formation of nucleolar R-loops. (**A**) Analysis of acrocentric chromosome fraction between 42WT and 42R in metaphase spreads from each cell line; t-test ****. (**B**) Representative images of endogenous nucleolin (NCL) immunofluorescence (IF). (**C**) Quantification of nucleolar area from images in (**B**) (**D**) Quantification of nucleolar circularity from images in B. IF with the R-loop binding S9.6 antibody in (**E**) 42WT, 72 hour TMZ treatment in 42WT, 42R, T98G. (**F**) Quantification of R-loop intensity in e. Scale bar (**B**) 50 µm (**E**) 5 µm. **, p<0.005, ****, p<0.0001.

Within nucleoli, RNA polymerase I (RNAPI) transcribes ribosomal RNAs (rRNA), which can hybridize to antisense ribosomal DNA (rDNA) to form rRNA:rDNA R-loops (23). We used the R-loop specific antibody S9.6 to visualize nucleolar R-loop changes (24). The TMZ-sensitive 42WT cells showed discrete, DNA damage-induced punctate R-loop accumulation throughout the nucleus (25) when treated with TMZ (Fig. 2E), compared to the nucleolar staining in DMSO- untreated 42WT cells (Fig. 2E). However, TMZ-resistant cell lines showed a robust increase in nucleolar R-loop accumulation, rather than discrete nucleoplasmic puncta associated with a normal DNA damage response as reported by Gorthi *et al* (25) (Fig. 2E, F). We validated the specificity of the R-loop antibody, S9.6, by treating cells with RNase H, which digests R-loops. This treatment abrogated specific nucleolar staining (Supplementary Fig. 1D). Overall, TMZ-resistant GBM cells exhibit a nucleolar stress response with an increase in the number of acrocentric chromosomes and nucleolar size, a decrease in nucleolar circularity, and differential R-loop staining patterns.

### Isoform usage changes in TMZ-resistant cells

WGS analysis further identified a significant increase in mutations in G-rich splice sites in TMZ-resistant cells (Fig. 3A). Splicing mutations and altered splicing patterns are characteristic of many cancers, including GBM (26). To quantify changes in categories of splicing events in the TMZ-resistant vs sensitive cells, we performed nanopore cDNA sequencing to interrogate full length transcripts. The Albacore workflow was used to call cDNA bases, followed by long-read alignment with minimap2 (27). Then both Isoform Switch Analyze R (28) and SUPPA2 (29) pipelines were used to identify changes in the differential gene isoform usage, between TMZ-S and -R cells, even in cases where there was no difference in overall gene expression (Fig. 3B). As other studies have investigated differentially expressed genes, we prioritized interrogation of genes with differential isoform usage. We first compared the switched gene expression between all three cell lines and found the highest overlap in switched gene expression between T98G vs 42WT and 42R vs 42WT, suggesting more common changes with resistance acquisition (Supplementary Fig 2A). We then analyzed the type of splicing changes and found differential exon number (EN) to be the top Isoform Switch Analyze R-annotated change (Fig. 3C). Next, we tested the occurrence of mutations within the isoform switched genes (121 of 153) as compared to non-switching genes (17, 815 of 32, 542), where Fisher’s Exact test showed a significant enrichment of mutations correlating to switched genes (p= 3.12e-10). Looking at the point mutation types in the switched genes; we again found the highest number of GàA and CàT mutations in these switched genes (Supplementary Fig. 2B). The group of 121 isoform-switch genes that also contained mutations was selected for further downstream analysis, where we first assessed global pathway changes via DAVID (30) to determine potential druggable vulnerabilities. We found an enrichment of genes in alternative splicing, protein binding, phosphorylation, acetylation, and the cytosol (Fig. 3D; Supplementary Table 1). We then interrogated functional changes of the switched isoforms and found a subset of genes that switched from nonsense-mediated decay (NMD)-sensitive to insensitive, or vice versa. For example, *FAM118A* switched from a dominant NMD-sensitive isoform to insensitive isoform, although overall gene expression shows decreased expression in resistant cells (Fig. 3E, Supplementary Fig. 2C). *FAM118A* has previously been identified as a potential target in glioma stem cells, in which a decrease in *FAM118A* gene expression, but increase in FAM118A protein levels, correlates with worse overall survival of patients with GBM(31). Conversely, *THUMPD2* shows an increase in total gene expression, but a switch to an NMD-sensitive isoform in the TMZ-resistant cells (Fig. 3F, Supplementary Fig. 2D). Decreased THUMPD2 protein expression is correlated with chemotherapy resistance (32). Overall, we show that TMZ-resistant cells have significant differential gene isoform usage as compared to TMZ-sensitive lines that correlates with gene mutation status.

**Figure 3.**
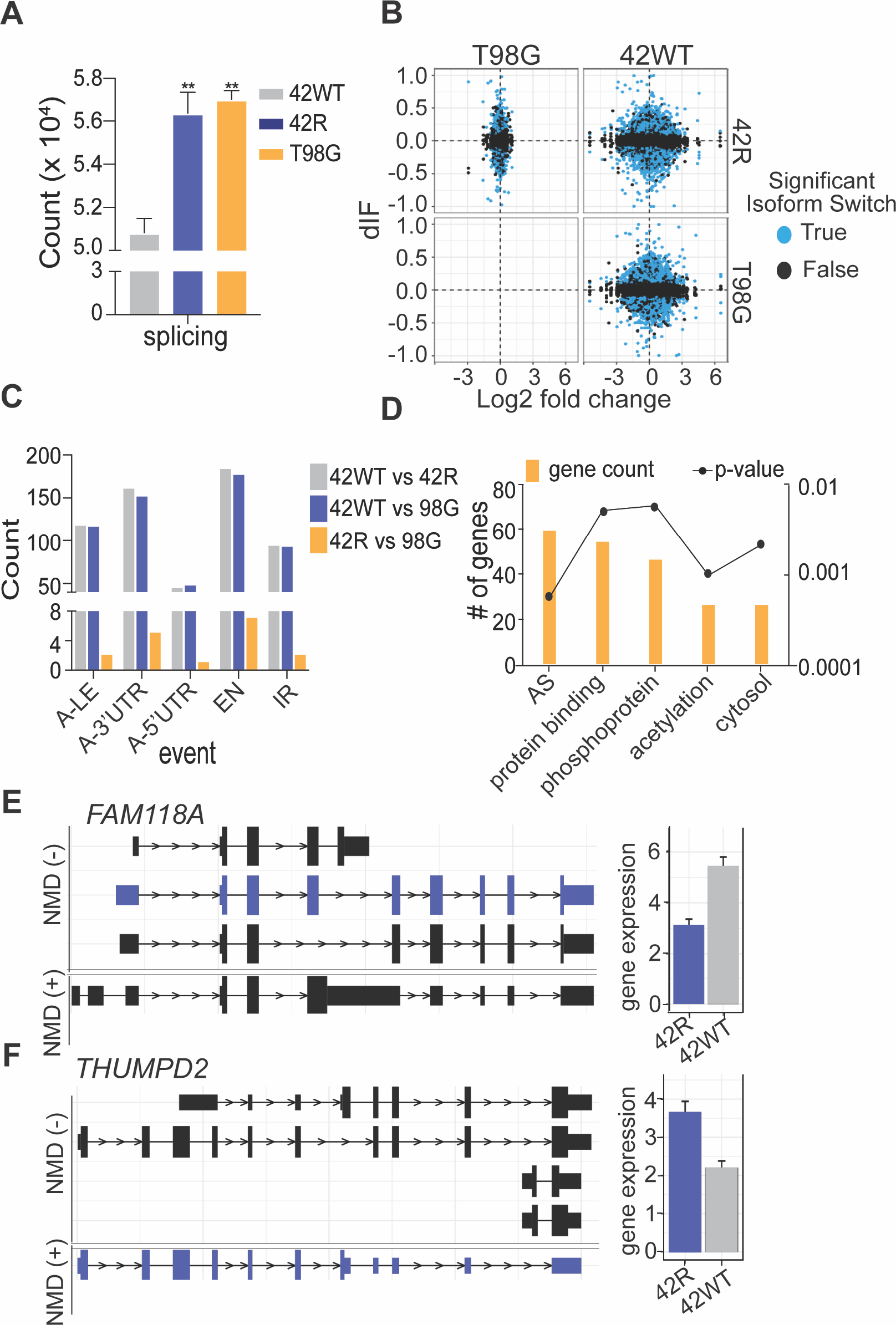
Splice-site mutations in TMZ-resistant GBM cells associate with increased exon and isoform usage changes in TMZ-resistant cells. (**A**) Quantification of splice-site mutations in TMZ-sensitive versus -resistant cells from whole genome sequencing analyses described in Fig. 1; One-way ANOVA. (**B**) Nanopore full-length cDNA sequencing analysis with Illumina short-read correction to determine the gene expression changes (x-axis) as compared to isoform usage changes (y-axis; dIF: differential isoform fraction). (**C**) Nanopore analysis of alternative splicing type, where the events are not mutually exclusive. Alternative 3’-UTR (A-3’UTR), Alternative 5’- UTR (A5’UTR), Alternative last exon (A-LE), Exon number change (EN), Intron retention (IR). (**D**) DAVID pathway analysis of gene subset which had both a gene isoform usage switch and annotated splicing mutation. Alternative splicing (AS). (**E**) *FAM118A* and (**F**) *THUMPD2* gene isoform changes between NMD-sensitive and -insensitive in TMZ-sensitive (42WT) vs TMZ-resistant (42R) cells.

### CLK2 inhibition preferentially targets TMZ-resistant cells

As our gene isoform usage analysis found significant changes in exon number and AS pathway regulation, we sought to functionally target RNA splicing in the TMZ-resistant cells. Several splicing factor families drive exon inclusion relative to exclusion events, including serine/arginine rich proteins (SRs). SR protein activity and localization are dependent upon their phosphorylation status, which can be altered by DNA damage (33). We found that the DNA-damaging TMZ treatment induced a significant decrease in the phosphorylation of SR proteins (pSRs) in TMZ-sensitive cells where DNA damage was measured by the presence of the double-strand break marker γH2AX (Fig. 4A). Dephosphorylation of pSRs was not observed in either TMZ-resistant cell line post TMZ-treatment (Fig. 4A), suggesting altered pSR function in TMZ-resistant cells. pSRs can bind both introns and exons to determine isoform fate through exon inclusion or exclusion events, and SR proteins that are not hyperphosphorylated are less competent to dictate gene isoform fate (34). Therefore, we sought to phenocopy the decrease in pSRs caused by TMZ treatment in the sensitive cells by inhibiting a pSR upstream regulator, the cdc2-like kinases (CLKs), in TMZ-resistant cells. CLKs catalyze the nuclear hyperphosphorylation of multiple SR proteins (35), with public data suggesting an association of higher expression of CLK2, but not CLK1, 3 or 4, with poor overall survival and increased grade in multiple brain cancers, including GBM (Fig. 4B, C). Consistent with this, our acquired and intrinsic TMZ-resistant GBM cells showed a robust increase in CLK2 protein expression as compared to the TMZ-sensitive line (Fig. 4D), but acute TMZ treatment of TMZ-sensitive cells did not increase CLK2 protein expression (Fig. 4D). GW807982X, here referred to as CLK2i, was published as part of the GSK Published Kinase Inhibitor Set (PKIS) (36) and identified as a CLK2 inhibitor. We further characterized this molecule in orthogonal assay systems and confirm potent and selective CLK inhibition and in cell target engagement. The compound is now a part of the Kinase Chemogenomic Set (KCGS) (37). Treatment of TMZ-resistant cells with CLK2i led to a decrease in SR phosphorylation at 30 minutes (Supplementary Fig. 3A-E) (38–40), and caused significant G2/M arrest, decreased cell proliferation, and increased apoptosis selectively in TMZ-resistant cells (Fig. 4E-G). Importantly, CLK2i treatment did not cause significant changes to growth rates or cell cycle, and only a modest increase in apoptosis, in the immortalized non-cancerous brain oligodendrocyte cell line, MO3.13 (Fig. 4E-G). These data support a model in which TMZ-resistant GBM cells have rewired their splicing regulatory machinery over time, as an increase in CLK2 expression was not observed following acute TMZ treatment (Fig. 4D). To further confirm increased CLK2 expression post TMZ-treatment, we re-analyzed patient samples from a cohort where three patients had matched RNAseq data from pre/post TMZ-treated tumors(41). In these three patients, all three recurrent GBM samples had increases of CLK2 expression, while the other CLK family members had varying changes in expression (Fig. 4H). Therefore, stably TMZ-resistant cells exhibit deregulation of SR protein phosphorylation and CLK2 function, and pharmacological inhibition of CLK2 may selectively target these alterations, leading to decreased cell proliferation and increased cell death.

**Figure 4.**
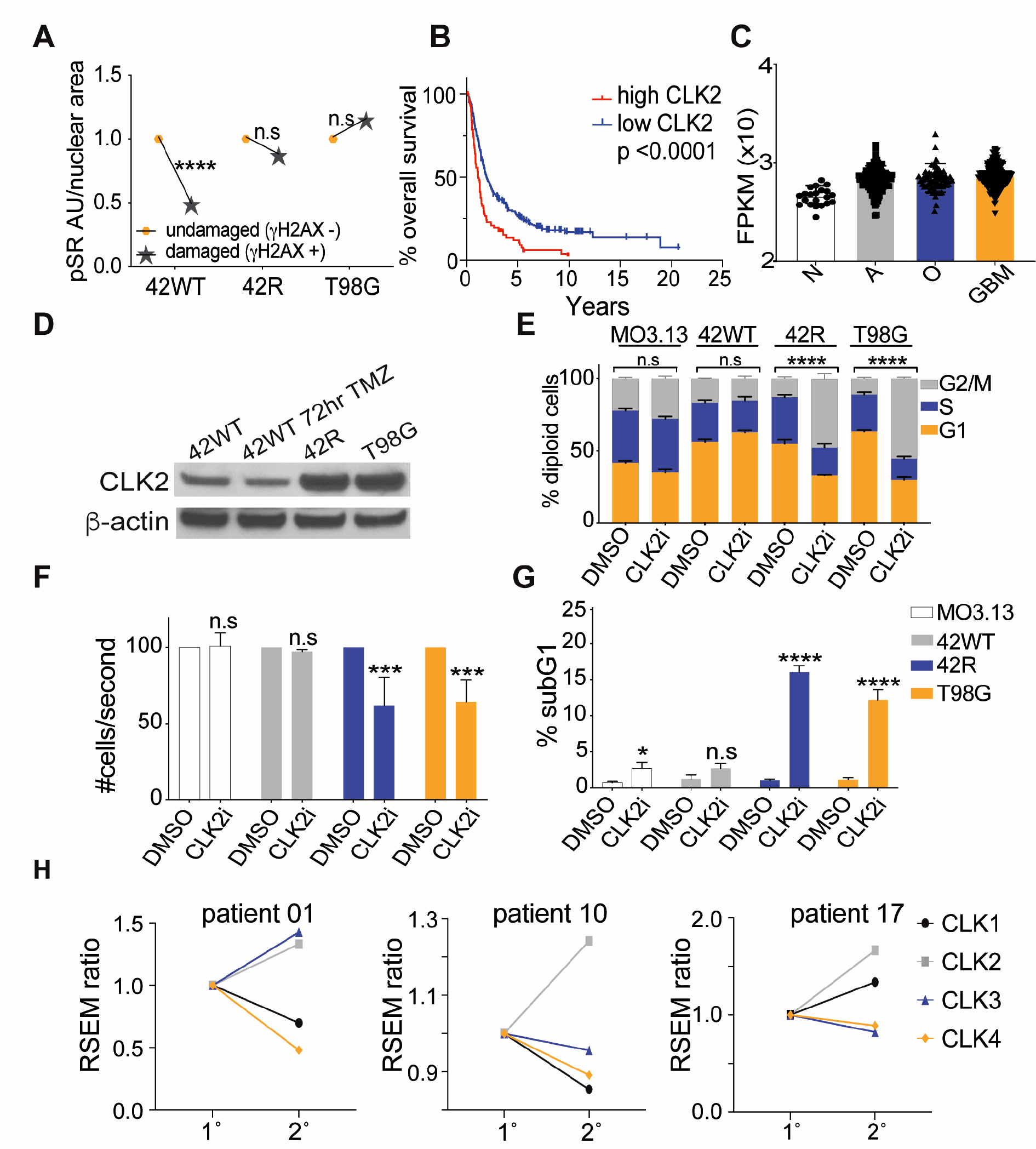
CLK2 inhibition with a novel CLK inhibitor (GW807982X, CLK2i) specifically targets TMZres GBM cells. (**A**) Quantified IF of pSR (1H4) intensity in TMZ-treated cells with (γH2AX+) or without (γH2AX-) DNA damage; two-tailed t-test. (**B**) Association between *CLK2* mRNA expression and overall survival in the REMBRANDT(70) brain cancer data set; log-rank (Mantel-Cox) test. (**C**) Association between *CLK2* expression and increasing tumor grade and aggressiveness in the REMBRANDT dataset; one-way ANOVA. Astrocytoma (A), Oligodendroglioma (O), glioblastoma (GBM). (**D**) Western blot of CLK2 expression in GBM TMZ-sensitive, -treated, and -resistant cell lines. (**E**) FACS analysis for cell cycle profile in response to 5 µM CLK2i vs. DMSO vehicle control treatment for 24 hour; three biological replicates with One-way ANOVA. (**F**) Relative cell numbers following 24 hour treatment with 5 µM CLK2i versus DMSO vehicle control; three biological replicates with t-test, (**G**) SubG1 or apoptotic DNA content in cells analyzed in g; three biological replicates with t-test. (**H**) Change in RNAseq RPKM of *CLK1-4* in 3 matched patient samples from primary (1°) GBM and post-TMZ treatment (2°)(41). * p<0.05, **p<0.001, ***p<0.0005, ****p<0.0001.

### EWSR1 cytoplasmic localization in TMZ-resistant cells and GBM clinical specimens

Given the collective deregulation of G4s, nucleoli, and splicing observed in TMZ-resistant GBM cells, we sought to identify a robust, independent marker that might be an indicator for TMZ resistance and, potentially, second line CLK2i efficacy in GBM. EWSR1 is member of the FET family of proteins, which also includes FUS and TAF15, where FUS is an established biomarker in many neurodegenerative disorders (13). EWSR1 binds DNA G4 structures (16), relocates to nucleoli following UV-induced DNA damage (42), and plays key functional roles in splicing and DNA damage responses (42). The nuclear staining of EWSR1 in TMZ-sensitive cells was in marked contrast to both the nuclear and cytoplasmic aggregates seen in acute TMZ-treated and TMZ-resistant cells (Fig. 5A, reversed image to better visualize the cytoplasmic staining, ). Importantly, expression levels of EWSR1 protein did not increase with TMZ resistance (Fig. 5B). The presence of cytoplasmic EWSR1 aggregates in TMZ-resistant cells was validated by a second antibody from a different vendor and host species (Supplementary Fig. 4A). We then used high resolution stimulated emission depletion (STED) microscopy to better resolve the structure of these aggregates, which showed discrete protein puncta within the cytoplasmic aggregates in both TMZ-resistant cell lines (Fig. 5C) and confirmed that the amyloid-like aggregates were only found in the cytoplasm, but not present in the nucleus (Supplementary Fig. 4B).

**Figure 5.**
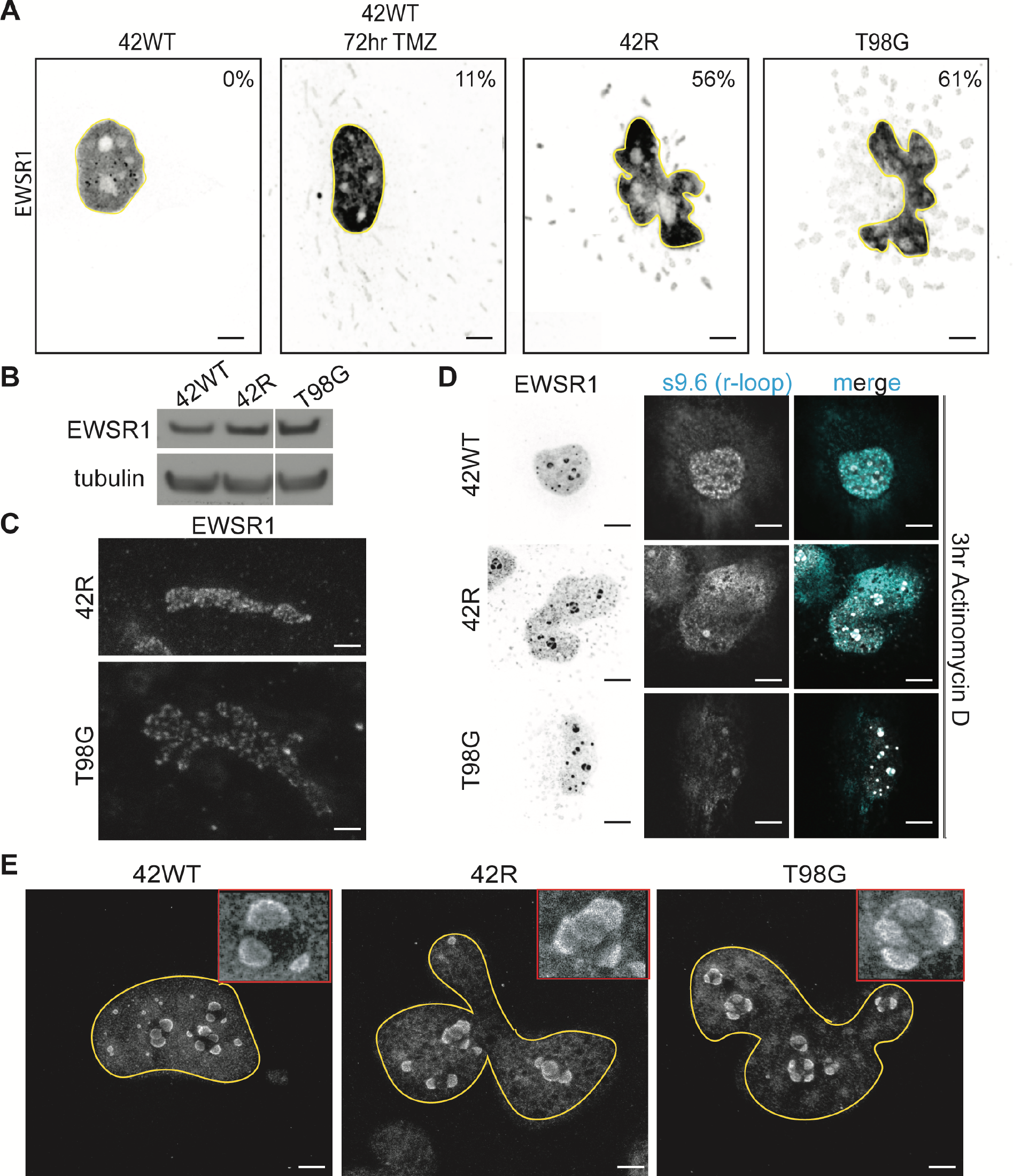
The RNA binding protein EWSR1 forms cytoplasmic, amyloid-like aggregates in TMZ-treated and TMZ-resistant GBM cells. (**A**) Basal IF of EWSR1 in the indicated cell lines, with the nucleus outlined in yellow. Values indicate percent of cells per condition that show EWSR1 cytoplasmic aggregates (**B**) Western blot of EWSR1 total protein expression in the indicates cell lines. (**C**) High-resolution stimulated emission depletion microscopy (STED) imaging of EWSR1 cytoplasmic aggregates in 42MBGA-TMZres T98G cells. (**D**) Localization of EWSR1 to nucleoli adjacent to R-loops (S9.6) following 3 hour 10 nM Actinomycin D (ActD) treatment to inhibit RNAPI. (**E**) STED imaging of EWSR1 nucleolar movement following 3 hour ActD treatment in 42WT 42R, and T98G cells. Scale bars in (**A**) 5 µm (**C**) 0.5 µm, (**D**) 5 µm, and (**E**) 2.5 µm.

Cytoplasmic aggregation of EWSR1 may suggest a potential role for RNA buffering, a defining feature of mis-localized RNA binding proteins in neurodegenerative diseases (43), where lower RNA concentration in the cytoplasm has been hypothesized to allow the aggregation of these proteins. As EWSR1 has already been shown to relocate to the nucleoli under stressed conditions (44), we sought to decrease the local RNA concentration and cause a stress response with low dose Actinomycin D (ActD) treatment, testing both an alternative (non-TMZ) stress response and the potential role of RNA in EWSR1 aggregation. Low dose ActD treatment led EWSR1 to accumulate around the nucleoli, indicated by s9.6 staining for R-loops (Fig. 5D). STED imaging of these samples better resolved EWSR1, with 42WT cells showing a horseshoe shaped staining pattern in which EWSR1-containing structures did not link to each other (Fig. 5E, left). By contrast, EWSR1 formed a linked structure between the droplets in both the acquired and intrinsic TMZ-resistant cells (Fig. 5E, middle and right). This further suggested a role for RNA buffering in EWSR1 aggregation. In addition, dual staining of EWSR1 and DNA G4s (BG4) showed colocalization in the TMZ-sensitive cell line that was not observed in TMZ-resistant cells (Supplementary Fig. 5), further suggesting that the deregulation of G4s identified in TMZ-resistant cells (Fig. 1C) can disrupt the localization of G4-binding proteins like EWSR1. However, EWSR1 aggregates do not appear to drive or maintain TMZ-resistance, as knockdown of EWSR1 had minimal effects on cell cycle profile or proliferation in TMZ-resistant cells (Supplementary Fig. 6A-C). We further used Leptomycin B (LMB), an inhibitor of the nuclear export protein chromosomal maintenance 1 (CRM1), to show that EWSR1 was efficiently trafficked to and from the cytoplasm in both TMZ-sensitive and -resistant lines (Supplementary Fig 6D). Ectopically expressed, YFP-tagged EWSR1 caused non-endogenous ribbon-like formation of EWSR1 in the cytoplasm in the TMZ-resistant cell lines, preventing further overexpression experiments (Supplementary Fig. 6E). Furthermore, neither EWSR1 nor CLK2 had significant isoform usage changes (Supplementary Table 1). Taken together, these results suggest that EWSR1 is the first, to our knowledge, aggregating RNA binding protein in GBM, where RNA buffering and deregulation of G4 structures may play a role in its aggregation.

**Figure 6.**
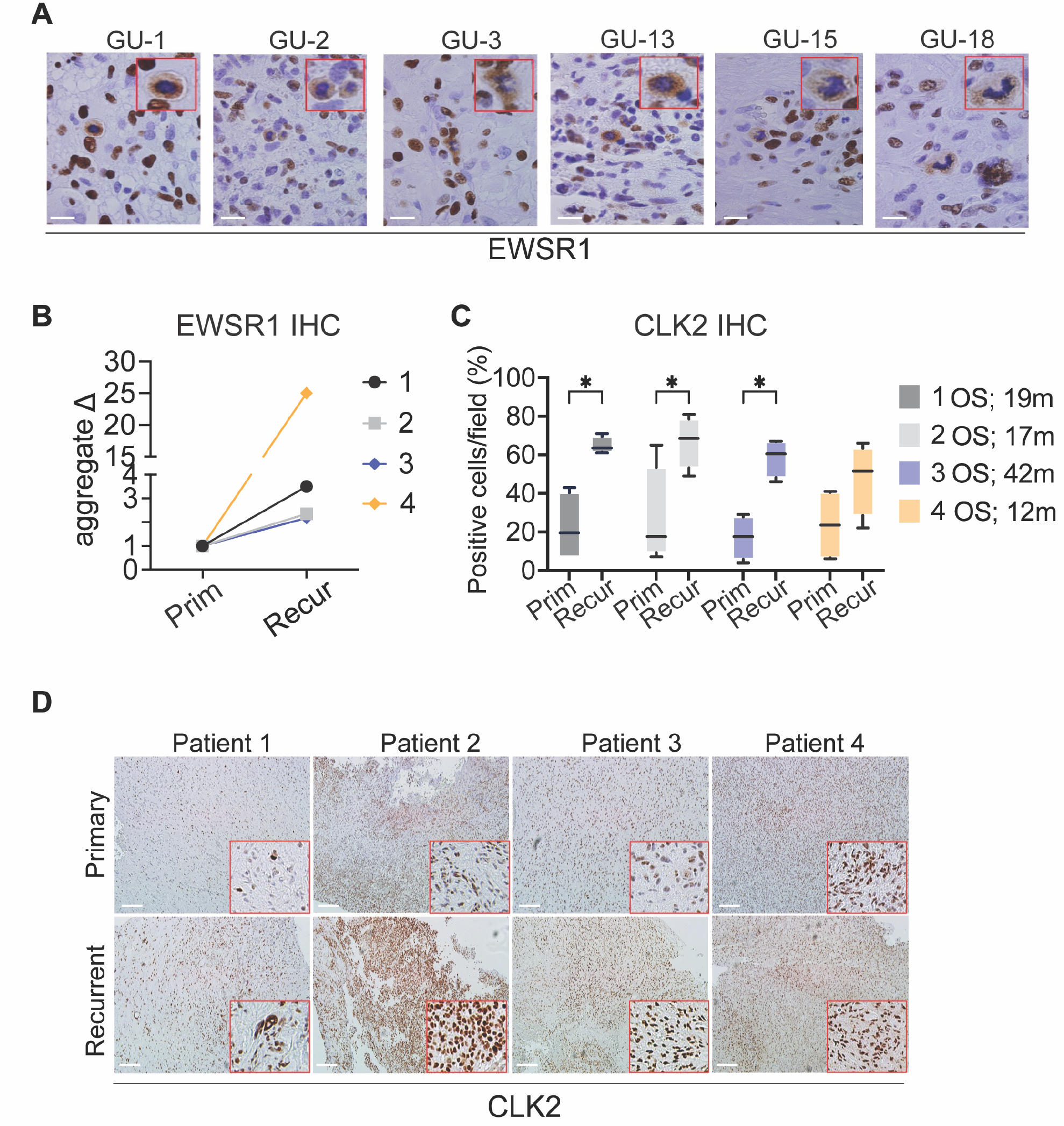
Cytoplasmic EWSR1 aggregates are detected in GBM clinical specimens. (A) EWSR1 IHC of 6 representative GBM patients with cytoplasmic aggregation of EWSR1 (red box inset). (**B**) Quantification of EWSR1 aggregative positive cells between Primary (Prim) and Recurrent (Recur) matched patient samples. (**C**) Quantification of CLK2 positive cells between Primary (Prim) and Recurrent (Recur) matched patient samples from images in (**D**) Overall survival (OS) in months (m). (**D**) Representative images of CLK2 IHC from matched patient samples. Scale bars in (**A**) 20 µm and (**D**) 100 µm *, p<0.05.

A key limitation of studies conducted in cell culture is their dependence on *ex vivo* models that have been in culture for decades. To determine whether EWSR1 cytoplasmic aggregates were also present in GBM clinical specimens, we stained a cohort of 21 brain tumor samples, including 15 GBMs, for EWSR1. Nine of the 15 GBM samples were positive for EWSR1 cytoplasmic amyloid-like aggregates (Fig. 6A, Supplementary Table 2), while none of the anaplastic astrocytomas or medulloblastomas showed this staining pattern. To further test our hypothesis that TMZ-resistant disease has an increase of EWSR1 cytoplasmic aggregates and CLK2 expression, we performed IHC on an independent set of 4 matched patient samples from pre-TMZ treatment to post-TMZ resistant recurrent GBM. We found an increase in EWSR1 cytoplasmic staining in all 4 samples, with the greatest relative increase observed the patient with shortest overall survival (OS, Fig. 6B). We also observed a significant increase in CLK2 expression in 3 of the 4 samples of post-TMZ resistant recurrent disease, with the fourth sample showing a high baseline CLK2 expression and the lowest overall survival (OS, Fig. 6C, D). Overall, this data establishes a clinically relevant framework for future studies to determine the functional role of EWSR1 aggregation and aberrant splicing via CLK2 protein abundance in GBM.

## Discussion

Since the pivotal 2005 clinical trial which changed GBM standard of care to the Stupp regimen - adjuvant TMZ post-surgery and radiation - TMZ has been almost exclusively used in brain cancers, despite adding only ∼4 months to median overall survival (45). Given the lack of successful FDA-approved second-line treatments for GBM, it is essential to identify pathways that could be targeted in TMZ-resistant GBM (46). However, TMZ is rarely used in other cancers, and the 400+ clinical trials to-date that have attempted to repurpose chemotherapeutic or targeted agents from other cancers have failed to demonstrate benefit of these approaches in GBM(47). We therefore looked to other central nervous system (CNS) disorders for insight. Mechanistic studies in CNS disorders, including neurodegenerative diseases (NDs), show pathological disruption of DNA and RNA secondary structures (48), splicing, and aberrant localization of RNA binding proteins (49). We therefore reframed our strategy for addressing TMZ-resistance in the context of nucleotide secondary structure, RNA processing, and RNA binding proteins in TMZ-resistant GBM.

As the first anticancer drugs were DNA-targeting, and still play a major role in cancer treatments, we focused on the downstream effects of TMZ-induced DNA damage at guanine nucleotides (8). Guanine is the most readily oxidized base, with the loss of an electron creating a hole that is preferentially targeted by spontaneous DNA damage, an event whose impact is compounded in tracts of guanines (50, 51). The established mechanism of action of TMZ is the addition of a mutagenic O^6^-methyl adduct to guanines (5). Guanines are critical nucleotides that stabilize many essential DNA and RNA secondary structures and are the target of many other chemotherapeutic agents (6). It also has previously been shown that TMZ-treated GBM has a characteristic mutational signature of C>T (18). By performing whole genome sequencing (WGS), we found both C>T and G>A mutations to be enriched in TMZ-resistant GBM cells. We focused on two critical G-rich regions – G-quadruplexes (G4s) and splice sites – to assess their functional roles in TMZ-resistance. We show sequence- and conformation-based changes in G4s by WGS and immunofluorescence, respectively, in TMZ-sensitive versus acute TMZ treated and TMZ-resistant GBM cells. We also demonstrate that the G4-stabilizing ligand TMPyP4 is more growth-inhibitory in TMZ-resistant lines compared to that in TMZ-sensitive cells. Previous studies have demonstrated the efficacy of other G4 ligands in multiple cancer cell lines, with CNS-derived models being the most sensitive to these agents (52). Therapeutic targeting of G4s in ALS can also decrease toxic hexanucleotide repeat foci and dinucleotide repeat proteins, leading to attenuated disease phenotypes *in vitro* (53). This avenue of targeting nucleotide secondary structure changes post-chemotherapy warrants further investigation, and will require the development of interventions that can cross the blood-brain barrier (BBB), as TMPyP4 cannot (54).

Our results reveal that TMZ-resistant GBM cells show an increase in a second critical DNA:RNA secondary structure: nucleolar R-loops. Nucleoli are major stress organelles, and aberrations in their size and circularity are positively correlated with increasing cancer grade (10). The increase in nucleolar R-loops we demonstrate in TMZ-resistant cells is intriguing, as a normal DNA-damage response would result in punctate R-loops throughout the nucleus (25), which we observed in TMZ-sensitive cells in response to TMZ. However, this pattern is lost in TMZ-resistant cells. In other settings, increases in nucleolar R-loops have been shown to induce genome instability (55), correlate with rDNA mutations, and an aberrant DNA damage response (23). This further suggests that there may be a reprogrammed DNA damage response in TMZ-sensitive versus -resistant cells.

We further focused on G-rich splice sites and their potential contribution to aberrant regulation of alternative splicing, a hallmark of many cancers including GBM (56). We showed that splicing mutations are enriched in the TMZ-resistant cell lines, where nanopore full-length cDNA sequencing also identified marked enrichment of changes in exon usage. The CLK family of splicing regulatory kinases induces the nuclear hyperphosphorylation of SR proteins. We show that SR proteins are de-phosphorylated upon TMZ treatment in TMZ-sensitive, but not -resistant, cells, there is an increase in CLK2 expression in TMZ-resistant cells, and a novel CLK2 kinase inhibitor is more growth-inhibitory in TMZ-resistant lines. CLK2 has previously been pharmacologically targeted in MYC-driven breast cancer (12) and genetically in GBM (57). Targeting splicing broadly is an active area of research with clinical trials underway in *SF3B1* mutant cancers and neurological diseases (58). Our data suggest that these efforts should be further expanded to TMZ-resistant GBM.

Our data identify EWSR1 as the first, to our knowledge, aggregating RNA-binding protein in TMZ-resistant GBM, using two independent models of acquired and intrinsic TMZ-resistant cell lines and GBM clinical samples. This aggregation phenotype is similar to DNA damaged-induced FUS aggregation observed in FUS-ALS motor neurons (59). RNA buffering may also play a role in the aggregation phenotype, as we show that inhibition of nucleolar rRNA transcription by acute low dose Actinomycin D (ActD) treatment can induce ribbon-like strands of EWSR1 around the nucleolus, though the local concentration of EWSR1 would presumably be increasing as well. The last factor we explored is the ability of DNA secondary structures to alter protein localization. The inability of EWSR1 to bind the mutated G4s in the TMZ-resistant lines suggest this may affect the normal function and proper regulation of EWSR1 in both splicing and DNA repair. Taken together, our data suggest potential overlapping functions of the cytoplasmic aggregates of EWSR1 with other neurodegenerative diseases, and further studies may provide important insight on specific upstream regulators of DNA damage-induced EWSR1 aggregation in CNS disorders and possibly other DNA damaging chemotherapy-treated cancers. However, significantly more research needs to be conducted to determine what role DNA and RNA secondary structures, RNA processing, and RNA binding protein aggregates play in GBM maintenance and TMZ-resistance.

In conclusion, this study illustrates a novel relationship between TMZ resistance and altered DNA/RNA structures, and how these changes can affect RNA binding protein function and localization. It further provides evidence that these changes may serve as therapeutic vulnerabilities that are targetable in TMZ-resistant glioma. Future research should be conducted to discover whether these results are generalizable to other chemotherapy-resistant cancers.

## Materials and Methods

### Cell Lines and Culturing Conditions

Immortalized human oligodendrocyte MO3.13 cells were a kind gift from Dr. Alexandra Taraboletti (Lombardi Comprehensive Cancer Center, LCCC). Patient-derived Temozolomide (TMZ) sensitive 42MGBA (42WT) cells were provided by Dr. Jeffrey Toretsky (LCCC), and the *de novo* TMZ resistant T98G cell line was from ATCC. The acquired TMZ resistant 42R cell line variant was developed by our lab and previously described(17). All cells tested negative for *Mycoplasma* contamination and were maintained in a humidified incubator with 95% air: 5% carbon dioxide. All cell lines were fingerprinted by the LCCC Tissue Culture and Biobanking Shared Resource (TCBSR) to verify their authenticity using the PowerPlex® 16 HS System (Promega, USA). The 42R cells are documented to be of the same origin as their parental cell line. MO3.13, 42WT, 42R, and T98G cells were grown in Dulbecco’s Modified Eagle Medium (DMEM, high glucose, ThermoFisher, #11965092) with 10% FBS.

### Immunofluorescence

Cells were seeded at a density of 25, 000-30, 000 cells onto 18 mm diameter #1.5 round coverslips (VWR, #101413-518) in 12-well dishes (day1). They were allowed to attach to the coverslips for a full day (day2). On the following day (day3), the media was removed, cells were washed 3x with PBS, and then fixed and permeabilized in 3.2% paraformaldehyde (PFA) with 0.2% Triton X-100 in PBS for 5 minutes at room temperature. Three washes were performed with PBS in the 12-well plate, then coverslips were inverted onto 120 μL of primary antibody in the antibody block (0.1% gelatin with 10% normal donkey serum in distilled H2O) on strips of parafilm and incubated for two hours. Coverslips were first incubated with either BG4 (Sigma MABE1126; 1:150), EWSR1 (Rabbit mAb Abcam ab133288; 1:600; Mouse mAb SCBT sc-48404; 1:200), S9.6 (Sigma MABE1095; 1:150), or NCL (Novus NBP2-44612-0.02mg; 1:500) for 2 hours. The donkey serum, non-confluent cells, 2-hour incubation for primary antibodies, and day3 staining is *vital* for visualizing consistent EWSR1 cytoplasmic staining. After incubation with primary antibodies, coverslips were washed three times with PBS. Then coverslips were inverted onto 100 µL of antibody block with secondary antibodies (Alexa Fluor 488 anti-mouse - 1:200, Life Technologies #A11029; Alexa Fluor 594 anti-rabbit – 1:200, Life Technologies A11037) and DAPI (DNA, 1:500 dilution) for 20 minutes in the dark. Coverslips were again washed 3x with PBS, then gently dipped four times into molecular biology-grade water before inversion onto one drop of Fluoro-Gel (with TES Buffer, Electron Microscopy Sciences, #17985-30) then allowed to air-dry in the dark for at least 10 minutes. Slides were stored at 4°C until image collection on the LCCC Microscopy & Imaging Shared Resource (MISR) Leica SP8 microscope with the 63X oil objective at 1.52 magnification.

### DNase I and RNase H treatment

Cells were seeded at a density of 25, 000-30, 000 cells onto 18 mm diameter #1.5 round coverslips (VWR, #101413-518) in 12-well dishes (day1). They were allowed to attach to the coverslips for a full day (day2). On the following day (day3), the media was removed, cells were washed 3x with PBS, and then fixed and permeabilized in 3.2% paraformaldehyde (PFA) with 0.2% Triton X-100 in PBS for 3 minutes at room temperature. 10 µL of a 1000 U of DNase I stock was added to 500 µL of DNase I buffer on cells for 10 minutes or 2 µL of a 5000 U of RNase H stock (NEB M0297S) was added to 500 µL of RNase H buffer on cells for 5 minutes at 45 °C. Cells were washed 3x with PBS and 3.2% PFA with 0.2% Triton X-100 was added again for 5 minutes post-DNase I treatment or RNase H treatment. Immunofluorescent staining was completed as stated above.

### Evaluation of acrocentric chromosome copy number changes induced by TMZ resistance

As previously described, metaphase spreads from 42WT and 42R cells were prepared using a standard protocol (17). Chromosomes were stained with 4’, 6’-diamindino-2-phenyloindole (DAPI) to determine total chromosome copy number and acrocentric chromosome copy number in each metaphase. Ratios of total chromosome number to acrocentric chromosome number in each metaphase were calculated and graphed in R.

### NanoBRET measurements for GW807982X (CLK2i)

CLK1, 2, and 4 NanoBRET assays were performed as previously described (60, 61). In brief the *N*-terminal Nano Luciferase/CLK1 fusion (NL-CLK1) or C-terminal Nano Luciferase/CLK2 fusion (CLK2-NL) or C-terminal Nano Luciferase/CLK4 fusion (CLK4-NL) was encoded in pFN31K expression vector, including flexible Gly-Ser-Ser-Gly linkers between NL and CLK1, 2 or 4 (Promega Madison, WI, USA) (62). For cellular NanoBRET Target Engagement experiments, the NL-CLK1 or CLK2-NL or CLK4-NL fusion construct was diluted with carrier DNA - pGEM- 3Zf (-) (Promega, Madison, WI, USA) at a mass ratio of 1:10 (mass/mass), prior to adding FuGENE HD (Promega, Madison, WI, USA). DNA:FuGENE complexes were formed at a ratio of 1:3 (μg DNA/μL FuGENE HD) according to the manufacturer’s protocol (Promega, Madison, WI, USA). The resulting transfection complex (1 part, volume) was then gently mixed with 20 parts (v/v) of HEK-293 cells (ATCC) suspended at a density of 2 x 10^5^ cells/mL in DMEM (Gibco) + 10% FBS (Seradigm/VWR). 100 μL of this solution was added to each well of a 96-well plate (Corning 3917) followed by incubation (37 °C/ 5 % CO2) for 24 hours. After 24 hours the media was removed from the HEK293 CLK (NL-CLK1, CLK2-NL or CLK4-NL) transfected cells and replaced with 85 μL OPTI-MEM media (Gibco). NanoBRET Tracer 5 (Promega, Madison, WI, USA) was used at a final concentration of 1.0 μM as previously evaluated in a titration experiment. A total of 5 μL/well (20x working stock of nanoBRET Tracer 5 [20 μM]) was added to all wells, except the “no tracer” control wells to which 5 μL/well of tracer dilution buffer alone was added. All inhibitors were prepared initially as concentrated stock solutions in 100% DMSO (Sigma). A total of 10 μL/well of the 10x chemical inhibitor stock solutions (final assay concentration 1 % DMSO) were added. For “no compound” and “no tracer” control wells, a total of 10 μL /well of Opti-MEM plus DMSO (9 μL was added (final concentration 1 % DMSO)). 96 well plates containing cells with NanoBRET Tracer 5 and inhibitors (100 μL total volume per well) were equilibrated (37 °C/ 5 % CO2) for 2 hours. To measure NanoBRET signal, NanoBRET NanoGlo substrate at a ratio of 1:166 to Opti-MEM media in combination with extracellular NanoLuc Inhibitor diluted 1:500 (10 μL [30 mM stock] per 5 mL Opti-MEM plus substrate) were combined to create a 3x stock. A total of 50 μL of the 3x substrate/extracellular NanoLuc inhibitor were added to each well. The plates were read within 15 minutes (GloMax Discover luminometer, Promega, Madison, WI, USA) equipped with 450 nM BP filter (donor) and 600 nM LP filter (acceptor), using 0.3 s integration time instrument utilizing the “nanoBRET 618” protocol. Eleven concentrations of GW807982X were evaluated in competition with NanoBRET Tracer 5 in HEK293 cells transiently expressing CLK1, 2, and 4. Prior to curve fitting the values are converted to mBRET units (x1000). Additional normalization of the NanoBRET assay data was performed by converting experimental values for respective concentrations of experimental inhibitors to relative percent control values (no compound [Opti-MEM + DMSO + Tracer 5 only] wells = 100 % Control, no tracer [Opti-MEM + DMSO] wells = 0 % Control). The data was normalized to 0 % and 100 % inhibition control values and fitted to a four-parameter dose-response binding curve in GraphPad Software (version 7, La Jolla, CA, USA).

### Cell cycle analysis

For TMPyP4 treatment, on day1 cells were seeded at 100, 000 cells per well in 6-well plastic tissue culture dishes one day prior to treatment with the indicated concentrations of drug. 50 µM TMPyP4 was added on day2, for an additional 48 hours. After 48 hours, cells were collected, washed with PBS, ethanol-fixed, stained with propidium iodide, and analyzed for cell subG1 (fragmented/apoptotic) DNA content and cell cycle profile. 20, 000 cells were acquired by flow cytometry on a Becton Dickinson Fortessa in the Flow Cytometry and Cell Sorting Shared Resource (FCSR). Files were modeled using ModFit software (Verity Software, Topsham, ME) to determine subG1, G1, S, and G2/M cell cycle stage. For CLK2i treatment, on day1 cells were seeded at 200, 000 cells per well in 6-well plastic tissue culture dishes one day prior to treatment with the indicated concentrations of drug. 5 µM CLK2i was added on day2, for an additional 24 hours. After 24 hours, cells were collected, washed with PBS, ethanol-fixed, stained with propidium iodide, and analyzed for cell subG1 (fragmented/apoptotic) DNA content, cell cycle profile, and growth rate. 20, 000 cells were acquired by flow cytometry on a Becton Dickinson Fortessa. Files were modeled using ModFit software (Verity Software, Topsham, ME) to determine subG1, G1, S, and G2/M cell cycle stage.

### Nucleolar size and circularity detection and graphing

Images taken from the Leica SP8 microscope were opened in Fiji is Just ImageJ (FIJI) (63), split into the channel to be analyzed, and converted from RGB to 8-bit images for analysis. In FIJI, the intensity threshold was set via Image -> Adjust -> Threshold (over/under 95). Next, the image was made into a binary form (Process -> Binary -> Make binary). Finally, particles that met the threshold were quantified for area and circularity via Analyze -> Analyze particles. These readouts were then imported into R and graphed/analyzed using the R package Raincloud.

### Western blot analysis

Cells were lysed in RIPA buffer supplemented with protease and phosphatase inhibitors (Roche, #4906837001) for protein extractions and separated by polyacrylamide gel electrophoresis using 4-12% gradient gels (Novex by Life Tech, #NP0321BOX) as described previously (19). They were then transferred onto Nitrocellulose membranes (Invitrogen, #IB23001) with the iBlot2 (Invitrogen, #IB21001) and probed with the following antibodies: EWSR1 (1:1000, abcam, ab133288), CLK2 (1:1000, sigma, HPA055366-100UL), phosphorylated SR proteins (clone 1H4, 1:500, Millipore, #MABE50. Beta-Tubulin (1:5000, Sigma Aldrich, #T7816) and Beta-Actin (1:5000, Sigma-Aldrich, #A5316) were used as loading controls. Proteins were detected with horseradish peroxidase-conjugated secondary antibodies (1:5000, GE Healthcare Life Sciences, #NA931-1ML (Mouse) or #NA934-1ML (Rabbit)) and enhanced chemiluminescent detection HyGLO Quick Spray Chemiluminescent (Denville Scientific, #E2400) using film (Denville Scientific, #E3212).

### High resolution stimulated emission depletion microscopy (STED)

STED images were acquired using a Leica SP8 3X STED microscope, a white-light laser for fluorescence excitation (470-670 nm), time-gated hybrid-PMTs, and a Leica 100x (1.4 N.A.) STED White objective (Leica Microsystems, Inc.). AF594 was excited with 575nm excitation, with a back projected pinhole at 200 nm, and the fluorescence emission was collected at 610 nm with a Line average or 4, Frame accumulation of 2, and Line accumulation of 1. Time-gating of the emission signal from the PMT was set to a range of 0.7-6.5 ns for experiments involving the 775 nm depletion laser. Z-stacks were taken X= 24.890 nm, Y=24.890 nm, and Z=160.217 nm. The pinhole was set to a value of 0.7 airy units for all images. Image deconvolution was performed using Hyugens software (Scientific Volume Imaging B.V., Netherlands) assuming an idealized STED point spread function.

### siRNA knockdown of EWSR1

Dharmacon individual set of 4 2 nmol EWSR1 siRNAs were (LQ-005119-02-0002) resuspended in RNA reconstitution buffer provided. On day 0, 250 µL of Opti-MEM was warmed and added to 5 µL of siRNA (stock = 20 µM) and 7.5 µL of Trans-IT X2. The mixture was left to form micelles for 25 minutes. This was then added dropwise to 6-wells with 1% FBS media and 100, 000 freshly plated cells (day1). The cells then adhered overnight, and media was changed the following morning. Cells were collected 48 hours later for either cell cycle analysis or Western blot, which allowed for 72 hours of transfection total.

### Immunohistochemistry staining (IHC) on human GBM patient samples

IHC was performed for EWSR1 and CLK2 in de-identified human GBM tumor samples (Supplementary Table 2). Five-micron sections from formalin fixed paraffin embedded tissues were de-paraffinized with xylenes and rehydrated through a graded alcohol series. Heat induced epitope retrieval (HIER) was performed by immersing the tissue sections at 98 °C for 20 minutes in 10 mM citrate buffer (pH 6.0) with 0.05% Tween. IHC was performed using a horseradish peroxidase labeled polymer from Agilent (K4003) according to manufacturer’s instructions. Briefly, slides were treated with 3% hydrogen peroxide and 10% normal goat serum for 10 minutes each and exposed to primary antibodies for EWSR1 (1/400, Abcam, ab133288) or CLK2 (1/800, Human Protein Atlas, HPA055366) overnight at 4 °C. Slides were exposed to the appropriate HRP labeled polymer for 30 min and DAB chromagen (Dako) for 5 minutes. Slides were counterstained with Hematoxylin (Fisher, Harris Modified Hematoxylin), blued in 1% ammonium hydroxide, dehydrated, and mounted with Acrymount. Consecutive sections with the primary antibody omitted were used as negative controls.

### RNA isolation

750, 000 of various cells were seeded on day1. The following day, cells were washed 3x with PBS, trypsinized, and pelleted into an Eppendorf tube. Then, 4 mL of TRIzol was added to each tube and incubated at room temperature for 5 minutes. 1 mL of each sample was aliquoted into 4 separate tubes. 200 µL of chloroform was added to each tube and vortexed briefly to mix. Samples incubated at room temperature for 5 minutes, were briefly vortexed, and then centrifuged at 12, 000 g for 10 min at 4 °C. 500 µL of the clear upper layer was collected from each tube and pooled into two 1 mL Eppendorf tubes where 500 µL of isopropanol was added and mixed thoroughly by multiple inversions. Samples were incubated for 15 minutes at room temperature, and then centrifuged for 15 minutes at 12, 000 g at 4 °C. Supernatant was then discarded, and pellet was washed with 750 µL 80% ethanol by multiple inversions. Sample was centrifuged for 5 minutes at 4 °C at 12, 000 g. Pellet was again washed with 80% ethanol by multiple inversions and centrifuged the same way as before. The supernatant was removed and discarded, and a second quick spin was done to remove any residual contaminating liquid. The RNA pellet was then air dried for 10 minutes before being resuspended in 100 µL of nuclease-free water. Total RNA was quantified using a NanoDrop 2000, and quality assurance was performed by Agilent BioAnalyzer.

### Illumina short-read sequencing

20 μg aliquots of total RNA were used for preparation of indexed, 150-base paired-end sequencing libraries using the TruSeq Stranded Total RNA Library Prep Kit (Illumina). Sequencing was performed on the Illumina NextSeq 550 instrument in the Genomics and Epigenomics Shared Resource (GESR) at a minimum of 65M reads per sample.

### Nanopore full-length cDNA sequencing

20 μg aliquots of total RNA were diluted in 100 μl of nuclease free water and poly-A selected using NEXTflex Poly(A) Beads (BIOO Scientific Cat#NOVA-512980). Resulting poly-A RNA was quantified and 50 ng aliquots were transferred to thin-walled Eppendorf PCR tubes. The biological poly-A RNA and a synthetic control (Lexogen SIRV Set 3, 0.5 ng) were prepared for cDNA synthesis and nanopore sequencing following the ONT SQK-PCS108 kit protocol with a few exceptions. During the reverse transcription step, Superscript IV was used, and the reverse transcription incubation time was increased to 15 minutes. After reverse transcription, PCR was performed using LongAmp Taq Master Mix (NEB) under the following conditions: 95°C for 30 seconds, 10 cycles (95°C for 15 seconds, 62°C for 15 seconds, 65°C for 15 minutes), 65°C for 15 minutes, hold at 4°C. The resulting cDNA libraries were quantified, and sequencing libraries were prepared using 700 ng of cDNA following the standard protocol for SQK-PCS108 (1D sequencing). Sequencing was performed on the GridION platform using ONT R9.4 flow cells and the standard MinKNOW protocol script (NC_48hr_sequencing_FLO-MIN106_SCK-PCS108).

### Nanopore full-length sequencing analysis with short-read Illumina correction

Albacore workflow (version 2.1.3) was used for base-calling cDNA data. Using minimap2 (27) with recommended parameters (-ax splice -uf -k14), reads were aligned to the GRCh38 (64) human genome reference in order to investigate possible novel isoforms. The latest GRCh38 assembly Similarly, short reads were aligned to the reference with spliced aligner HISAT2 (65). In order to utilize both the long and short read alignments together in assembly, StringTie (66) was used in “--mix” mode to identify and quantify isoforms some of which are novel (67). These novel and Analyze R (28).

The genes that were identified to have statistically significant changes in isoform expression levels between cell lines were filtered through 3 stages:

1. The dominant Open Reading Frame (ORF) of the isoforms were identified by Isoform Switch Analyze R and switches between isoforms that result in the same ORF sequence were filtered out.
2. To analyze the switching events that possibly correlate with TMZ resistance, only the genes that show switching in both 42R and T98G cell lines when compared to the 42WT cell line were included.
3. With only two replicates per cell line, the gene switch Q-values generated by Isoform Switch Analyze R cannot confidently assess biological variation. To eliminate any potential false positives due to low number of replicates, a secondary differential splicing analysis was conducted with SUPPA2 (a tool that exploits between-replicate variability and can accurately evaluate PSI values using two replicates per condition) (29). Stage (2) was also applied to SUPPA2 results, and only the overlapping switch genes between Isoform Switch Analyze R and SUPPA2 results were included.

The final list of genes after filtering was investigated for mutation enrichment (using the genome arithmetic framework of BEDTools (68)) and the downstream pathways that were potentially affected. Files and codes associated with this analysis can be found at https://github.com/timplab/Drug_resistant_GBM.

### DNA isolation for whole genome sequencing

750, 000 cells were collected for each cell line, where DNA was isolated using the DNeasy Blood and Tissue Kit (Qiagen; Cat. No. 69504). Briefly, cells were centrifuged and resuspended with proteinase K and buffer AL for 10 min at 56 °C. 200 µL of ethanol was then added and mixed thoroughly by vertexing. DNA was added to the DNeasy Mini spin column, centrifuged, and flow- through discarded. DNA was washed in the DNeasy column with buffer AW1 once, and AW2 once as well. Membrane was then dried by centrifugation with all flow-through being discarded. DNA was eluted using 200 µL buffer AE after 1 min incubation at room temperature.

### Whole genome sequencing using Illumina TruSeq DNA PCR-Free Library prep Kit

Illumina TruSeq DNA PCR-Free Library prep Kit was used per manufacture instructions by the Genomics and Epigenomics Shared Resource (GESR). Paired-end, indexed libraries for human whole genome sequencing were constructed from 1.0 µg gDNA using the TruSeq DNA PCR-Free Library Prep Kit (Illumina, San Diego) according to manufacturer’s instructions. Briefly, DNA was fragmented using a Covaris M220 focused ultrasonicator (Covaris, Woburn, MA) using settings for a 350 bp insert size. Library quality was assessed with a BioAnalyzer 2100 using the High Sensitivity DNA kit (Agilent Technologies, Santa Clara, CA). The libraries were quantified using the Kapa Library Quantification Kit Illumina Platforms (Kapa Biosystems, Boston, MA). The denatured and diluted libraries were sequenced on a NextSeq 550 System (Illumina) using v2.5 High Output 300 cycle kit with 1% PhiX to an average sequencing depth of 50X coverage.

### Whole genome sequencing analysis

Raw data were first demultiplexed into 3 groups, which were 42WT, 42R and T98G, and each group had two biological duplicates. All raw sequence data (fastq files) were run through fastQC v0.11.7 for quality check, lower quality reads (Q < 20) were eliminated. Cutadapt v2.1 was used for adapter trimming, read length less than 20 bp after trimming were filtered. Processed reads were then aligned to GRCh38 reference sequence using bwa v0.7.17 paired-end mode. Mutation detection was conducted in GATK v4.1.0.0(69) following the best practice for variant calling workflow. The consensus assembly for each sample were created by bcftools, then the Quadron methodology (19) was used to detect stable G4 structures. Unique G-Quad sequence with Q > 19 were extracted from each group for further analysis.

## Acknowledgements

We wish to thank Drs. Deborah Berry, Angela Brooks, Christian Combs, Karen Creswell, Brent Harris, Rong Hu, Peter Johnson, Moshe Levi, Supti Sen, Alexandra Taraboletti, Jeffrey Toretsky, Todd Waldman, Cameron Soulette and Dan Xun for sharing reagents, scientific insights, technical assistance, and/or editorial comments on the manuscript. Technical services were provided by the following LCCC Shared Resources that are supported in part by NIH P30 CA051008: Flow Cytometry and Cell Sorting Shared Resource (FCSR); Genomics and Epigenomcs Shared Resource (GESR); Histopathology and Tissue Shared Resource (HTSR); Microscopy and Imaging Shared Resource (MISR); Survey, Recruitment, and Biospecimen Collection Shared Resource (SRBSR); and Tissue Culture and Biobanking Shared Resource (TCBSR).

## Funding

This work was supported by: a Georgetown University Medical Center (GUMC) Dean for Research’s Toulmin Pilot Project Award, and a Partners in Research Breakthrough Award, and pilot funding from NIH P30 CA051008 (to RBR); NIH K00 CA234799, a student research grant from the Medical Center Graduate Student Organization, and fellowship support from NIH T32 CA009686 (to DMT); Intramural Research Program, National Heart, Lung, and Blood Institute, National Institutes of Health (to JRH); NIH NS115403 and the Malnati Brain Tumor Institute of Northwestern Medicine (to SYC).); NIH HG010538 (to WT); and The Caroline Fund (to SLG). The Structural Genomics Consortium (applicable to CAF, DHD, CIW, JEP, and WJZ) is a registered charity (no: 1097737) that receives funds from AbbVie; Bayer Pharma AG; Boehringer Ingelheim; Canada Foundation for Innovation; Eshelman Institute for Innovation; Genentech; Genome Canada through Ontario Genomics Institute (OGI-196); EU/EFPIA/OICR/McGill/KTH/Diamond Innovative Medicines Initiative 2 Joint Undertaking (EUbOPEN grant no: 875510); Janssen, Merck KGaA, Merck Sharp and Dohme, Novartis Pharma AG; Pfizer, São Paulo Research Foundation-FAPESP, Takeda and the Wellcome Trust. Additional funding for the SGC-UNC was provided by The Eshelman Institute for Innovation, UNC Lineberger Comprehensive Cancer Center, PharmAlliance, and National Institutes of Health (1R44TR001916-02, 1U24DK116204-01).

## Author Contribution Statement

DMT contributed to study design, performed experiments, analyzed data, and wrote the paper. BE and RR contributed to study design and analyzed data. LJ analyzed data. NS performed experiments. CAF performed experiments. JRH provided reagents and input on the paper. BRH performed experiments and provided input on the paper. DHD provided reagents and input on the paper. CIW and JEP performed experiments. XS, AG, and BH provided input on the paper and generated figures. SG provided reagents and input on the paper. WJZ, MP, and WT contributed to study design, provided reagents, and input on the paper. SYC contributed to study design and analyzed data. RBR contributed to study design, analyzed data, and development of the paper. All authors reviewed, edited, and approved the manuscript.

## Competing Interests

WT has two patents (8, 748, 091 and 8, 394, 584) licensed to Oxford Nanopore. All other authors declare no potential conflict of interest.

## Ethics Statement

Tumor samples were collected by the Georgetown University Histopathology and Tissue Shared Resource (HTSR) Human Tissue Bank (HTB) under IRB protocol 1992-048, the Survey, Recruitment, and Biospecimen Collection Shared Resource (SRBSR) under IRB protocol Pro00000007, or the John Theurer Cancer Center Tissue Biorepository under IRB protocols Pro2108-0589 and Pro2018-0020. All experimental methods are in compliance with the Helsinki Declaration.

## Data and materials availability

The whole genome DNA sequencing, nanopore full-length RNA sequencing, and Illumina short-read RNA sequencing raw data generated and analyzed in the current study are available from the corresponding authors upon request and are deposited as BioProject PRJNA780296

---

**Supplementary Figure 1.**
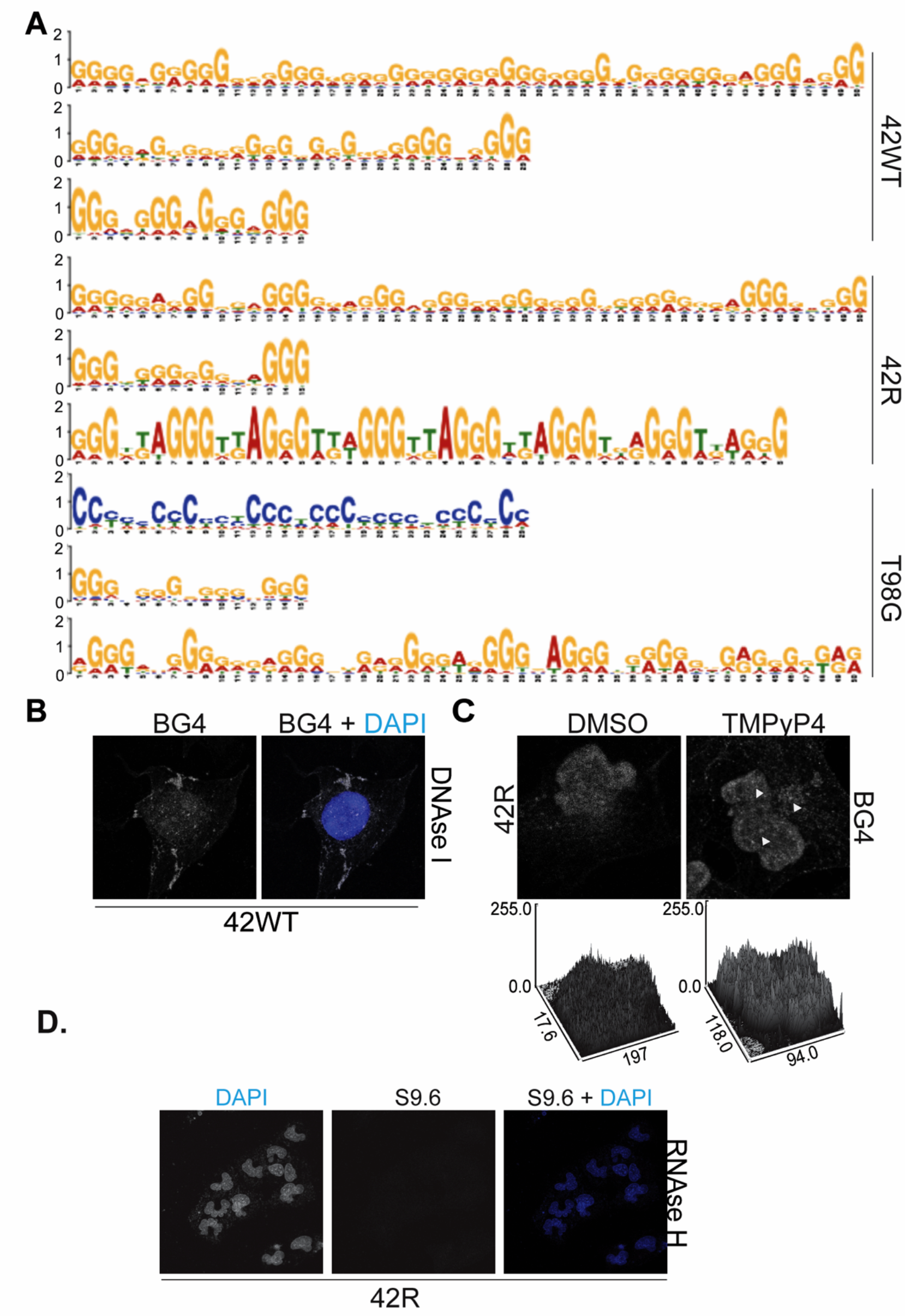
Specificity controls for BG4 and TMPyP4. (**A**) MEME-predicted enriched motifs from Quadron-derived unique acrocentric chromosome sequences between 42WT (E-value top; 1.4e-231, bottom; 9.4e-013), 42R (E-value 7.9e-206), and T98G (E-value top; 9.3e- 198, bottom; 7.2e-013) cell lines. (**B**) DNase I treatment of 42WT cells abrogates BG4 G-quadruplex antibody staining. (**C**) Treatment with 50 µM TMPyP4 vs. DMSO for 24 hour in 42R increases discrete punctate staining of BG4 and increased surface plot height. Nuclear surface plot of signal intensity depicted on the right was generated by FIJI. (**D**) RNase H treatment of 42R cells abrogates S9.6 R-loop antibody staining.

**Supplementary Figure 2.**
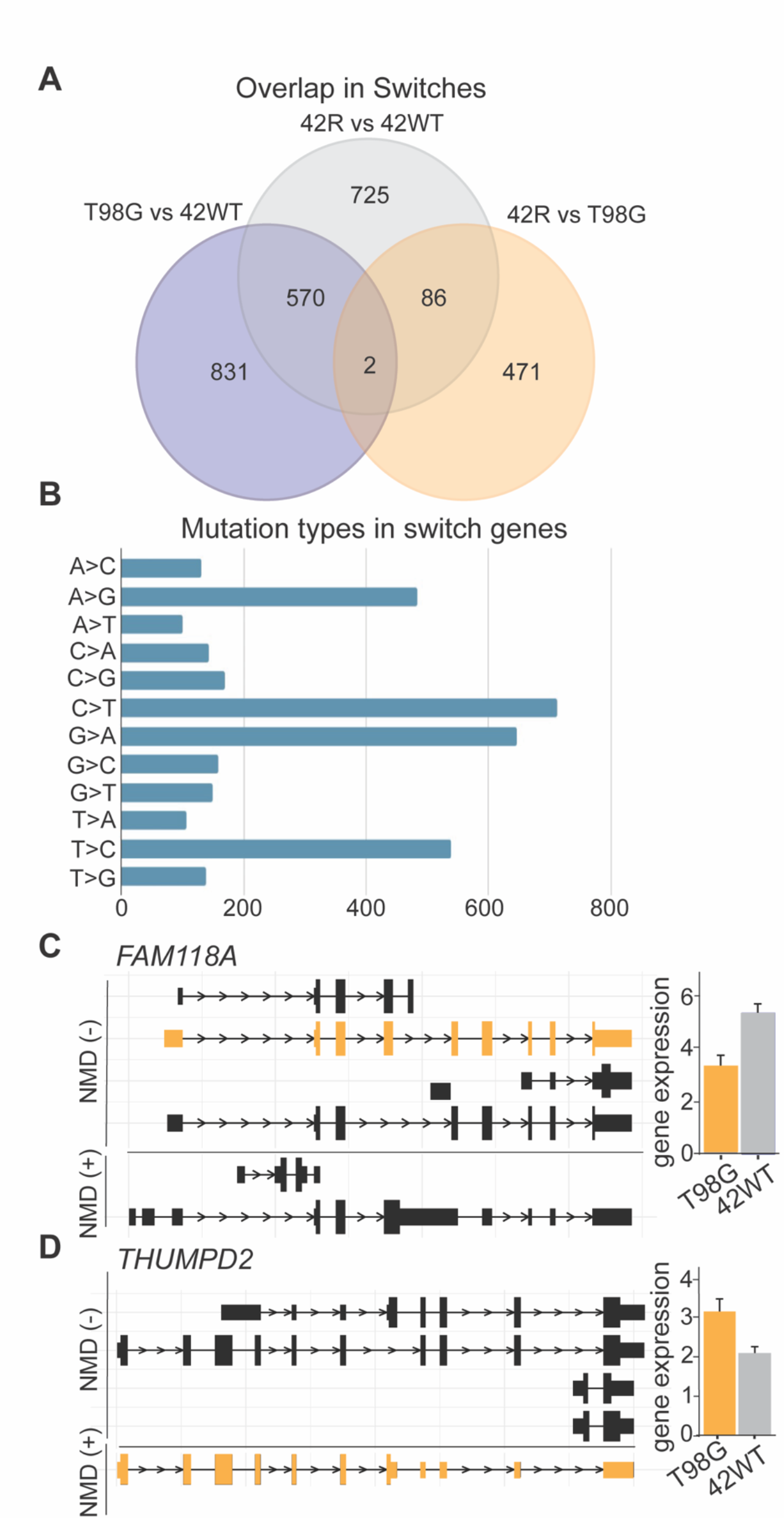
Additional nanopore and WGS mutation results. (**A**) Venn Diagram of the overlap in switched isoforms between the indicated cell lines. (**B**) Mutation type abundance in switched genes that also have splicing mutations. (**C**) *FAM118A* and (**D**) *THUMPD2* gene isoform changes between NMD-sensitive and -insensitive in TMZ-sensitive (42WT) versus TMZ-resistant (T98G) cells.

**Supplementary Figure 3.**
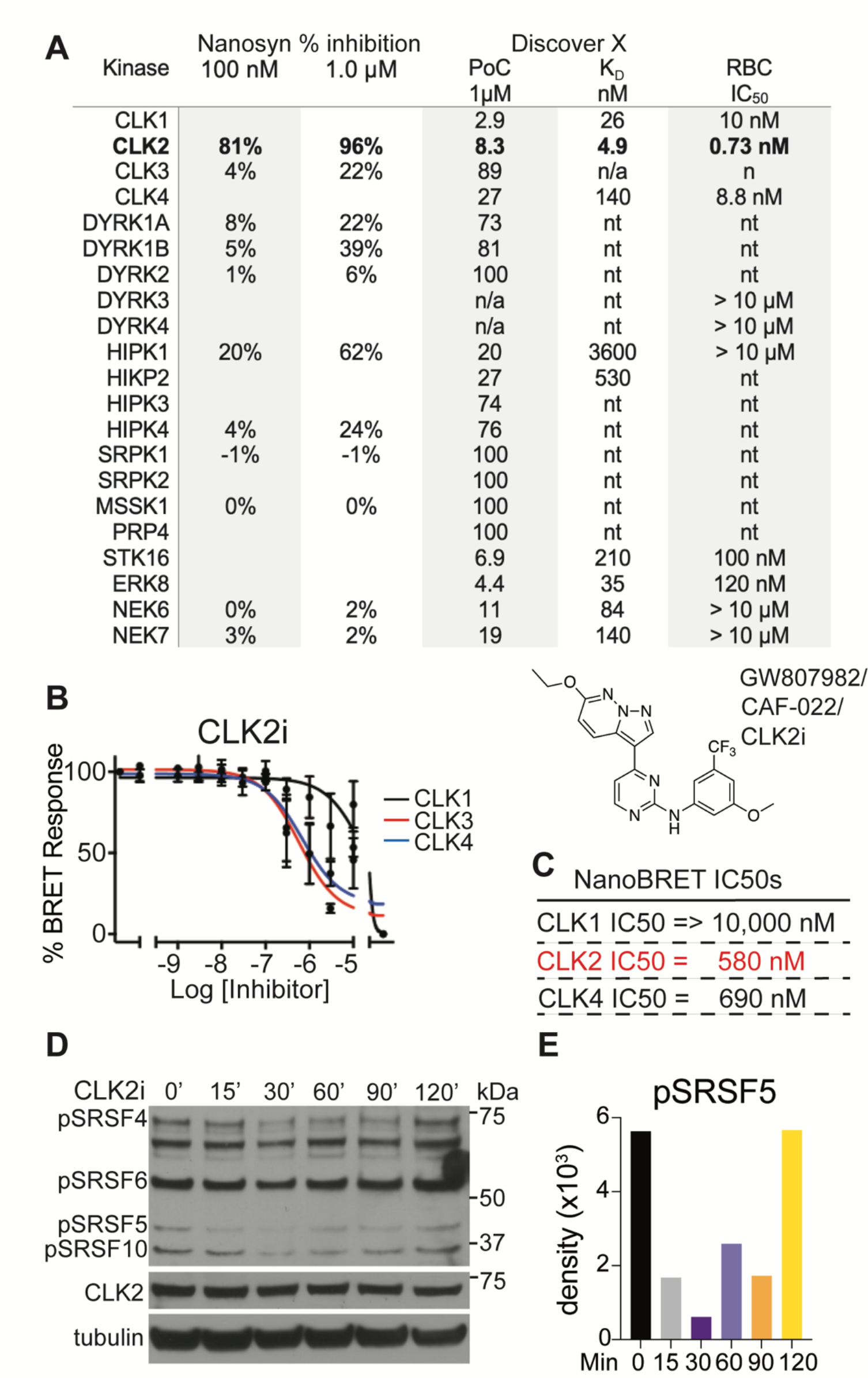
Characterization of a novel CLK2 inhibitor (GW807982X, CLK2i). (**A**) Commercial kinase binding and activity assays were performed at NanoSYN (data included are percent inhibition at 100 nM and 1 µM CLK2i; assays were conducted at [ATP] = *K*m^app^ of each kinase), DiscoverX (data included are *K*D values), and Reaction Biology Corporation (data expressed are IC50 values of kinase inhibition; assays were conducted at 10 µM ATP). “nt” denotes that the compound was not tested in these assays. (**B**) NanoBRET assay (77) of CLK2i (CAF-022) with multiple CLK family members in live cells, structure of CLK2i/CAF-022/GW907982X. (**C**) NanoBRET IC50 values determined for CLK1, 2, and 4. (**D**) Western blot of pSR changes over a time-course of CLK2i treatment in 42R cells (**E**) quantification of maximal pSRSF5 hypophosphorylation in (**D**). CLKi, GW807982X and CAF-022 are all names for the same compound. The NanoSYN data shown in (A) were originally published in (48)

**Supplementary Figure 4.**
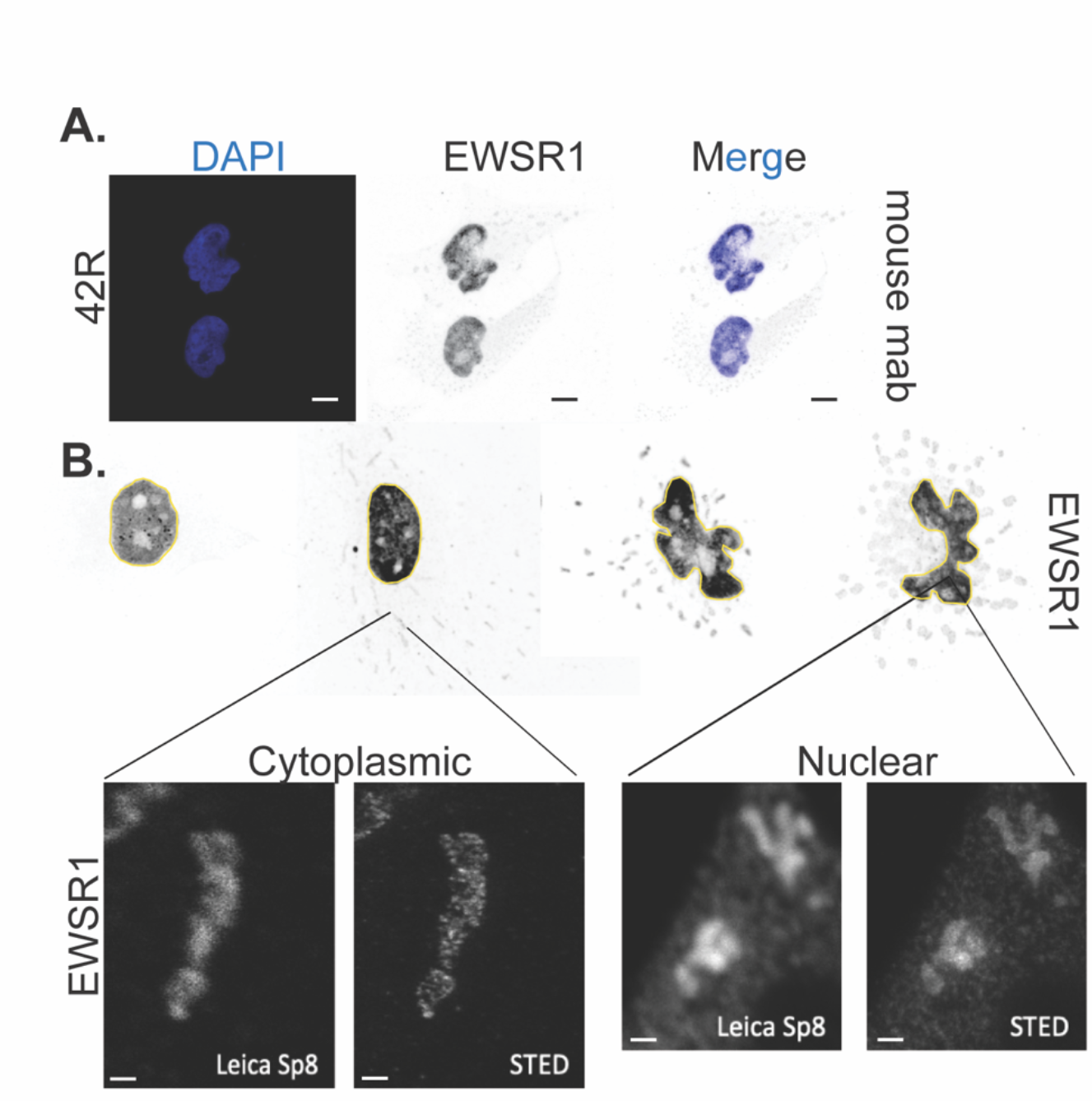
Controls for EWSR1 IF, aggregation, and STED imaging. (**A**) 42R cells stained with the EWSR1 mouse monoclonal antibody (mAb, Santa Cruz sc48404) and DAPI, scale bar 10 µm. (**B**) Staining with the EWSR1 rabbit mAb in 42WT, 42WT-treated with 100 µM TMZ for 72 hr, 42R, and T98G cells, higher magnification of nuclear EWSR1 staining is shown in both Leica Sp8 confocal and STED images, scale bar 2.5 µm.

**Supplementary Figure 5.**
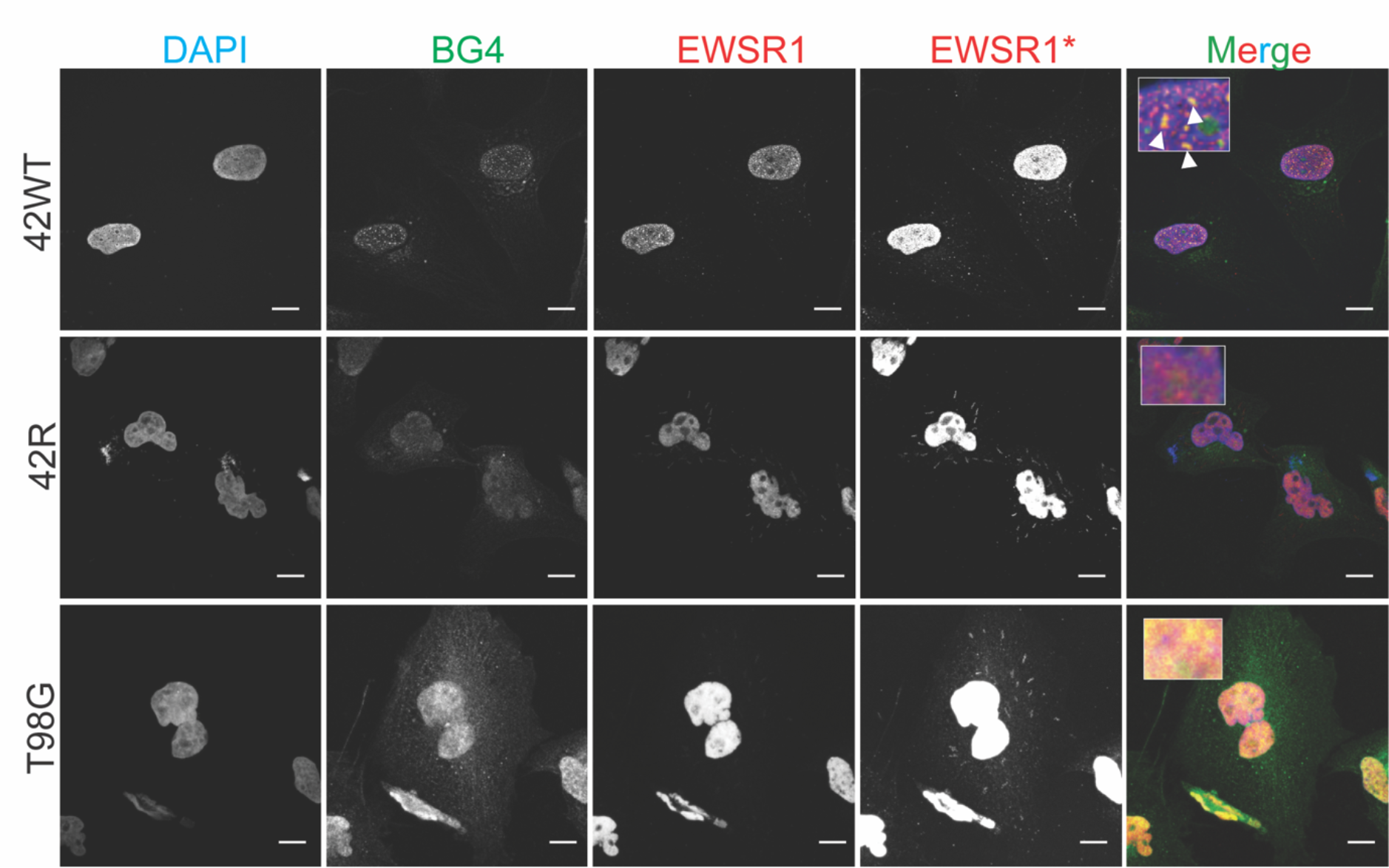
Controls for EWSR1 colocalization with G4s. Co-stain of the G4 antibody BG4 with EWSR1 in 42WT, 42R, T98G cells. EWSR1* shows enhanced images to permit visualization of cytoplasmic aggregates. Merged panels show an inset with higher magnification of BG4 and EWSR1 localization, arrows in 42WT show yellow colocalization of EWSR1 and BG4 staining. Scale bar 50 µm.

**Supplementary Figure 6.**
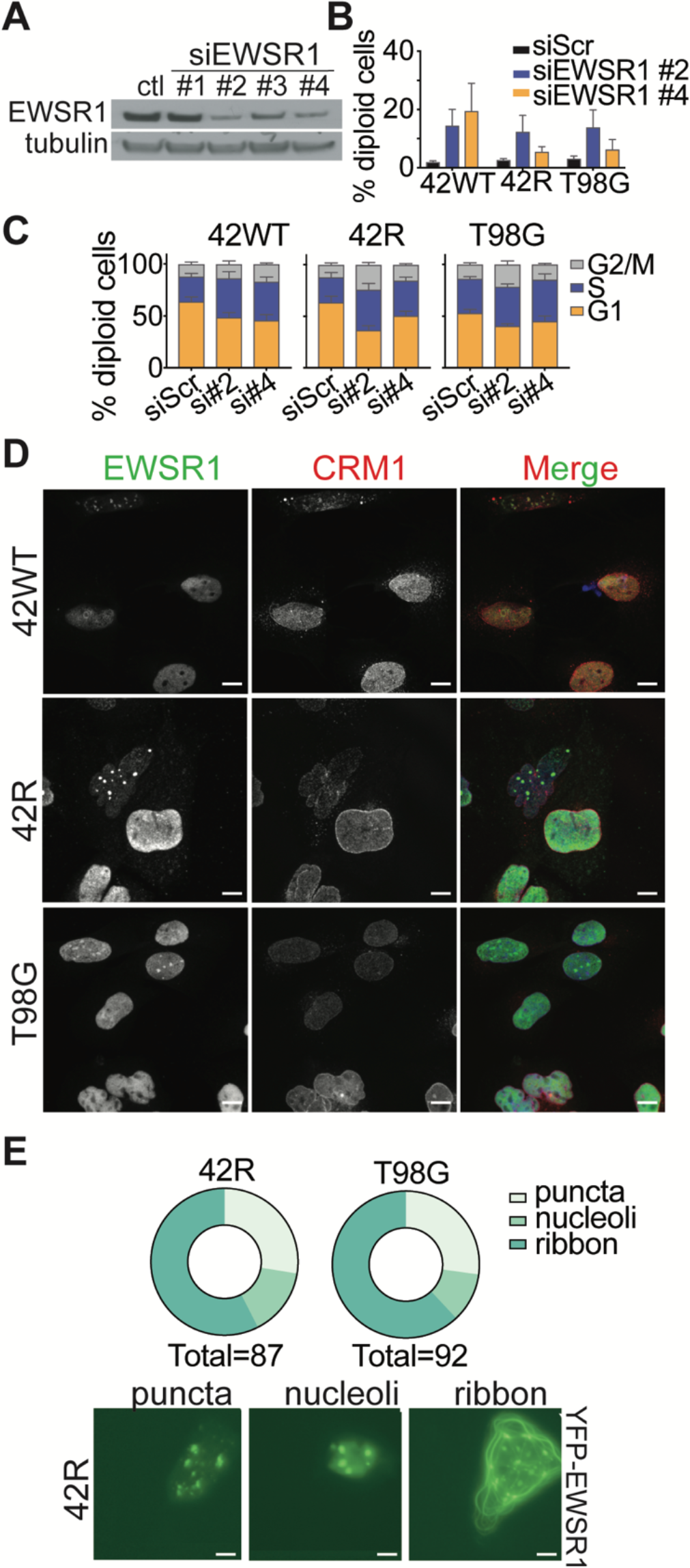
Controls for endogenous EWSR1 depletion and nucleo-cytoplasmic trafficking, and ectopic YFP-EWSR1 cytoplasmic aggregation. (**A**) Western blot of EWSR1 knockdown with four different siRNAs. (**B**) FACS analysis of subG1 DNA content following 72 hourof EWSR1 depletion. (**C**) Cell cycle profile of cells analyzed in (**B**) for 42WT, 42R, T98G cells; three biological replicates. (**D**) Treatment with 25nM Leptomycin B (CRM1 inhibitor; LMB) in 42WT, 42R, and T98G cells for 24 hr. (**E**) Quantification of the phenotype resulting from overexpression of YFP-EWSR1 phenotype in 42R n=87 cells, and T98G n=92 cells. Representative images below for “puncta”, “nucleoli”, and “ribbon” phenotypes. Scale bars in (**D)** 50 µm and (**E**) 10µm.

**Supplementary Table 1.**
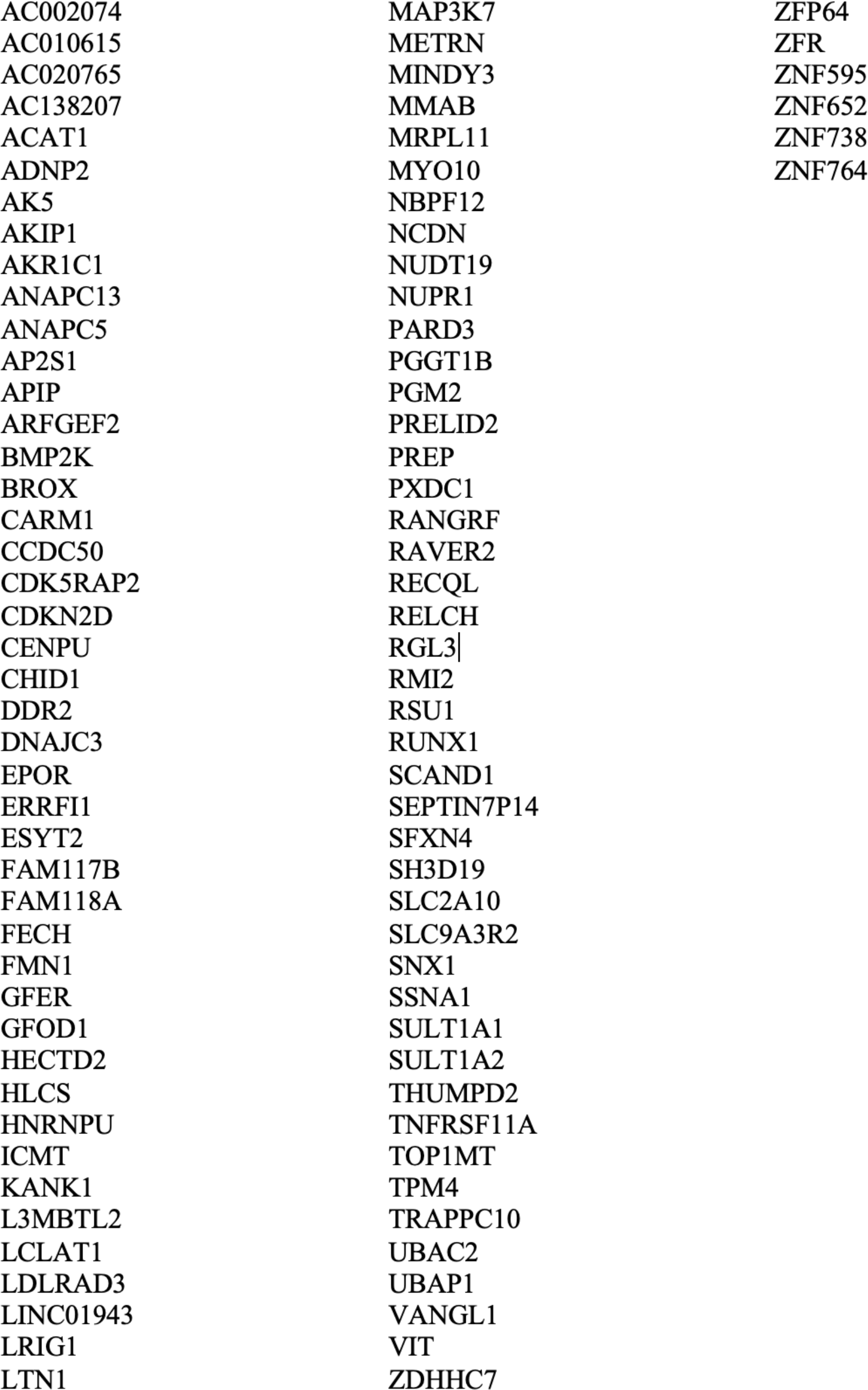
List of significant genes with both splice site mutations and isoform usage changes.

|  |  |  |
| --- | --- | --- |
| AC002074 | MAP3K7 | ZFP64 |
| AC010615 | METRN | ZFR |
| AC020765 | MINDY3 | ZNF595 |
| AC138207 | MMAB | ZNF652 |
| ACAT1 | MRPL11 | ZNF738 |
| ADNP2 | MYO10 | ZNF764 |
| AK5 | NBPF12 |  |
| AKIP1 | NCDN |  |
| AKR1C1 | NUDT19 |  |
| ANAPC13 | NUPR1 |  |
| ANAPC5 | PARD3 |  |
| AP2S1 | PGGT1B |  |
| APIP | PGM2 |  |
| ARFGEF2 | PRELID2 |  |
| BMP2K | PREP |  |
| BROX | PXDC1 |  |
| CARM1 | RANGRF |  |
| CCDC50 | RAVER2 |  |
| CDK5RAP2 | RECQL |  |
| CDKN2D | RELCH |  |
| CENPU | RGL3\| |  |
| CHID1 | RMI2 |  |
| DDR2 | RSU1 |  |
| DNAJC3 | RUNX1 |  |
| EPOR | SCAND1 |  |
| ERRFI1 | SEPTIN7P14 |  |
| ESYT2 | SFXN4 |  |
| FAM117B | SH3D19 |  |
| FAM118A | SLC2A10 |  |
| FECH | SLC9A3R2 |  |
| FMN1 | SNX1 |  |
| GFER | SSNA1 |  |
| GFOD1 | SULT1A1 |  |
| HECTD2 | SULT1A2 |  |
| HLCS | THUMPD2 |  |
| HNRNPU | TNFRSF11A |  |
| ICMT | TOP1MT |  |
| KANK1 | TPM4 |  |
| L3MBTL2 | TRAPPC10 |  |
| LCLAT1 | UBAC2 |  |
| LDLRAD3 | UBAP1 |  |
| LINC01943 | VANGL1 |  |
| LRIG1 | VIT |  |
| LTN1 | ZDHHC7 |  |

**Supplementary Table 2.**
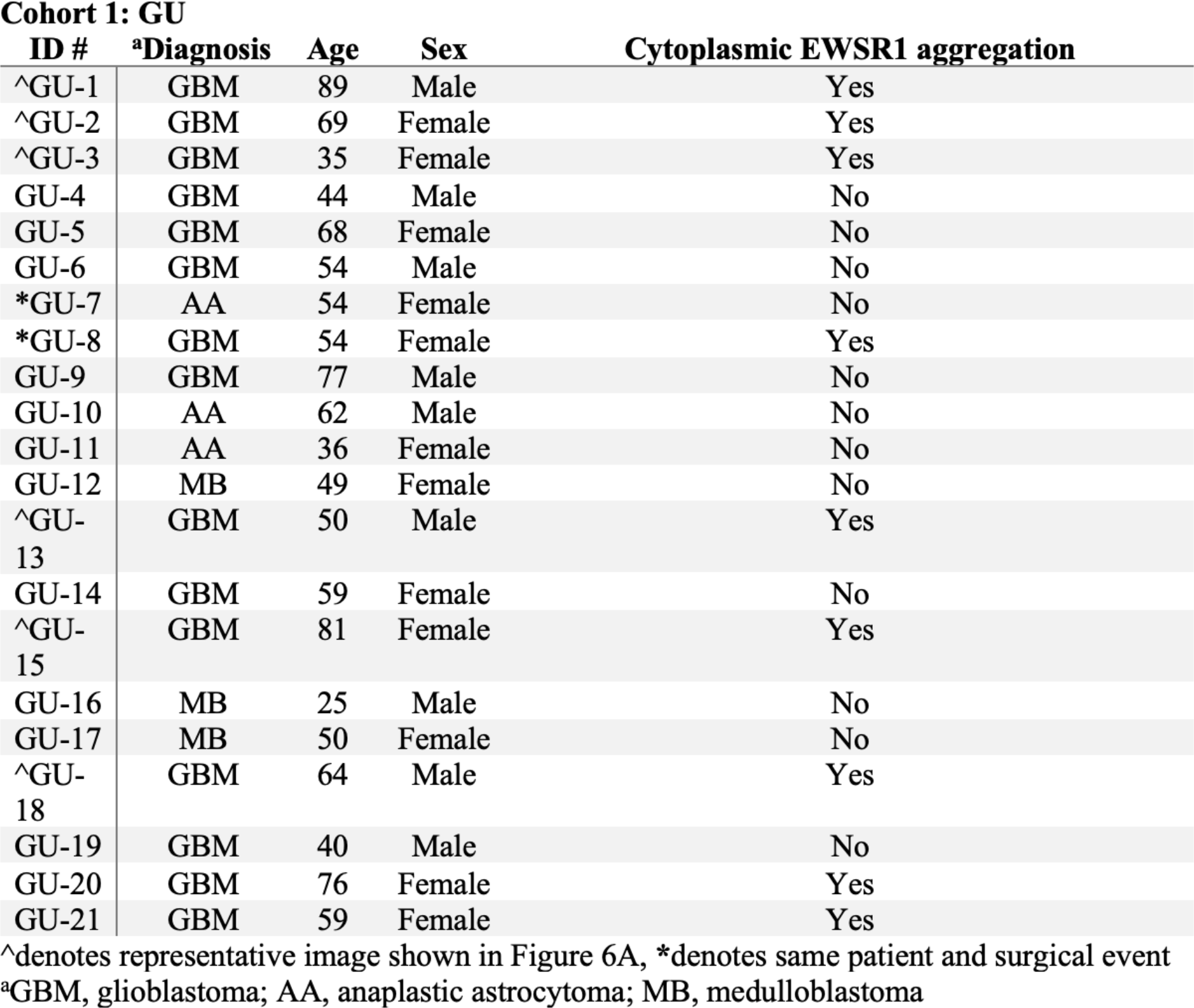
Demographic, clinical, and pathological characteristics of GBM clinical samples

## Notes

### Summary of Updates

Figures revised, new data and analyses.

